## Supplementary figures and tables for "Harmonious genetic combinations rewire regulatory networks and flip gene essentiality"

Supplementary Figures 1-8 and and Supplementary Tables 1-11

Aaron M. New and Ben Lehner

### Figure S1

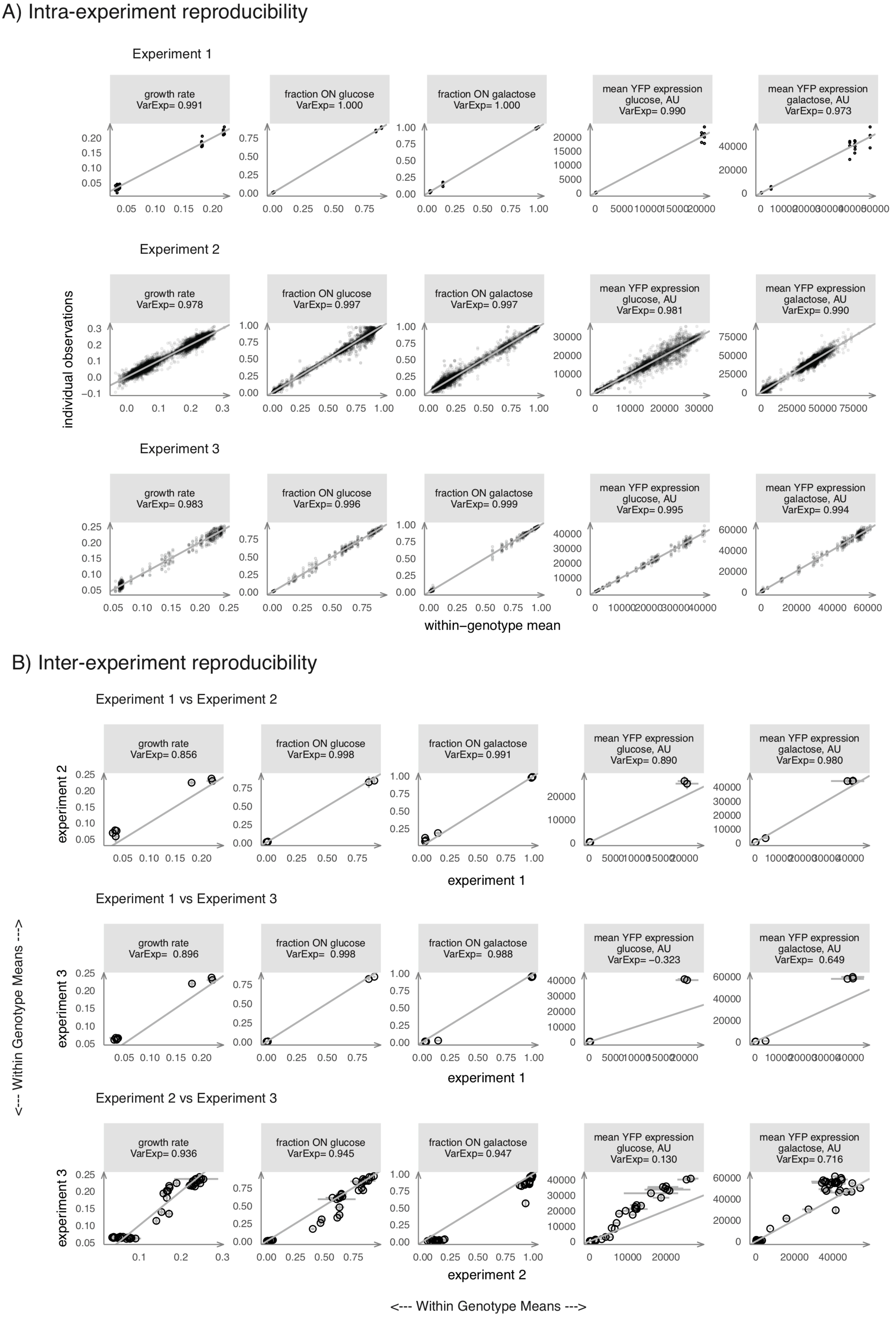
A) Intra-experiment reproducibility
Experiment 1
growth rate VarExp= 0.991
fraction ON glucose VarExp= 1.000
fraction ON galactose VarExp= 1.000
mean YFP expression glucose, AU VarExp= 0.990
mean YFP expression galactose, AU VarExp= 0.973
Experiment 2
growth rate VarExp= 0.978
fraction ON glucose VarExp= 0.997
fraction ON galactose VarExp= 0.997
mean YFP expression glucose, AU VarExp= 0.981
mean YFP expression galactose, AU VarExp= 0.990
Experiment 3
growth rate VarExp= 0.983
fraction ON glucose VarExp= 0.996
fraction ON galactose VarExp= 0.999
mean YFP expression glucose, AU VarExp= 0.995
mean YFP expression galactose, AU VarExp= 0.994
individual observations
within−genotype mean
B) Inter-experiment reproducibility
Experiment 1 vs Experiment 2
growth rate VarExp= 0.856
fraction ON glucose VarExp= 0.998
fraction ON galactose VarExp= 0.991
mean YFP expression glucose, AU VarExp= 0.890
mean YFP expression galactose, AU VarExp= 0.980
experiment 2
experiment 1
Experiment 1 vs Experiment 3
growth rate VarExp= 0.896
fraction ON glucose VarExp= 0.998
fraction ON galactose VarExp= 0.988
mean YFP expression glucose, AU VarExp= −0.323
mean YFP expression galactose, AU VarExp= 0.649
experiment 3
experiment 1
Experiment 2 vs Experiment 3
growth rate VarExp= 0.936
fraction ON glucose VarExp= 0.945
fraction ON galactose VarExp= 0.947
mean YFP expression glucose, AU VarExp= 0.130
mean YFP expression galactose, AU VarExp= 0.716
experiment 3
experiment 2
<--- Within Genotype Means --->
<--- Within Genotype Means --->

Supplementary Figure 1 continued

C

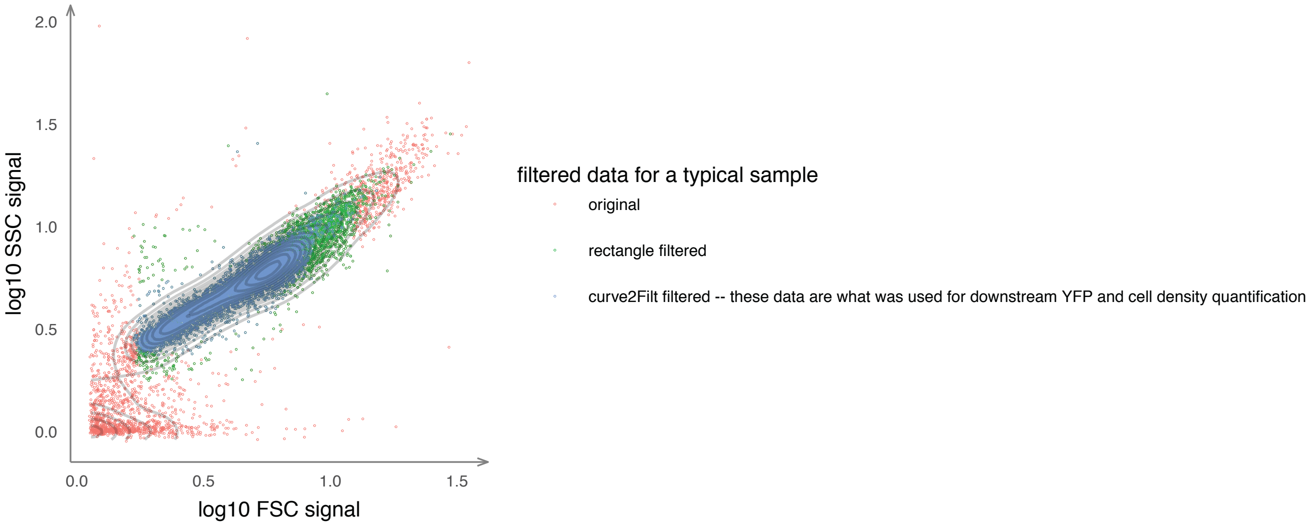

**Supplementary Figure 1: Reproducibility of the five key phenotypic values within and between experiments.** Panels are faceted by phenotype measurements (horizontally) and experiment or experiment comparison. The information above each panel includes the phenotype and the variance explained between the data underlying the two dimensions. The grey lines are where X = Y. a) The mean phenotype of each unique genotype in the dataset ('within-genotype mean'; horizontal axis) is plotted against each individual observation (vertical axis). b) Within-genotype mean values for a given experiment were paired with the within-genotype means of the same genotypes from the other two experiments and are plotted as circles. Error bars are the 95% confidence interval. c) A figure exemplifying the gating strategy based on cell size (FSC) and shape (SSC) features is illustrated. Two filtering steps, a rectangular and next a data-driven filter that selects high-density regions in two dimensions via kernel density estimation ("curve2filter") were used downstream of the original data (colours). A contour plot is shown over the 10000 data points to illustrate the overall distribution of a typical experiment's FSC and SSC values.

Supplementary Figure 2

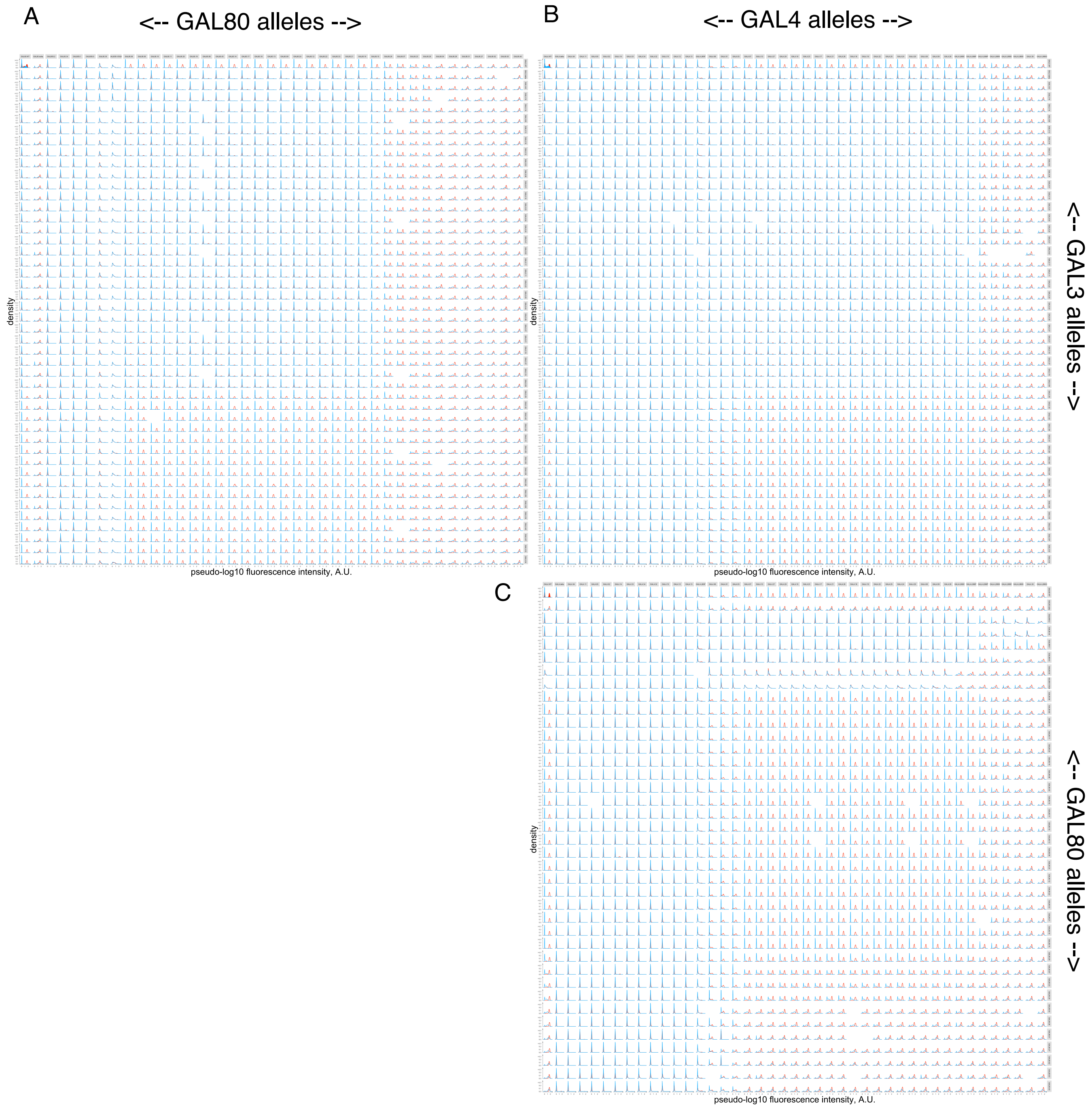

**Supplementary Figure 2: The distribution of Gal1p-YFP expression across all single and double mutants from the second pairwise mutant experiment.**

The figure is divided into three sections a-c, each with a GALR locus pairing (*GAL80* vs *GAL3*, *GAL3* vs *GAL4* and *GAL4* vs *GAL80*, respectively). The genotypes of the GALR allele are indicated in the strip text at the top or right-hand side of each section's panel sets. Each panel is a unique genotype. The single mutant allele ordering along the facetting dimensions were chosen first by calculating a 'phenotypic index' equal to fraction ON in glucose + fraction ON in galactose as a gross measure of how active the pathway was for a given clone. Following this statistic, roughly, a phenotypic index value of 0 = Uninducible, a value of 1 = Inducible and a value of 2 = Constitutive. Single alleles were then ordered according to the mean within-class phenotypic index value across single alleles within the pairing, followed by the mean within-genotype phenotypic index of single mutants. Finally, the WT and ∆ control strains were added to the top of the order in order to help the eye examine the single-mutant effects in a WT background and expectation of clone behavior when the other GALR locus was deleted. Lines and shading are the mean and ± 1 SD of genotypes falling into the given genotype (N=4 for N=2 independent transformations). Blue lines are samples growing in glucose and red lines are samples after 12 hours of growth in galactose.

### Supplementary Figure 3

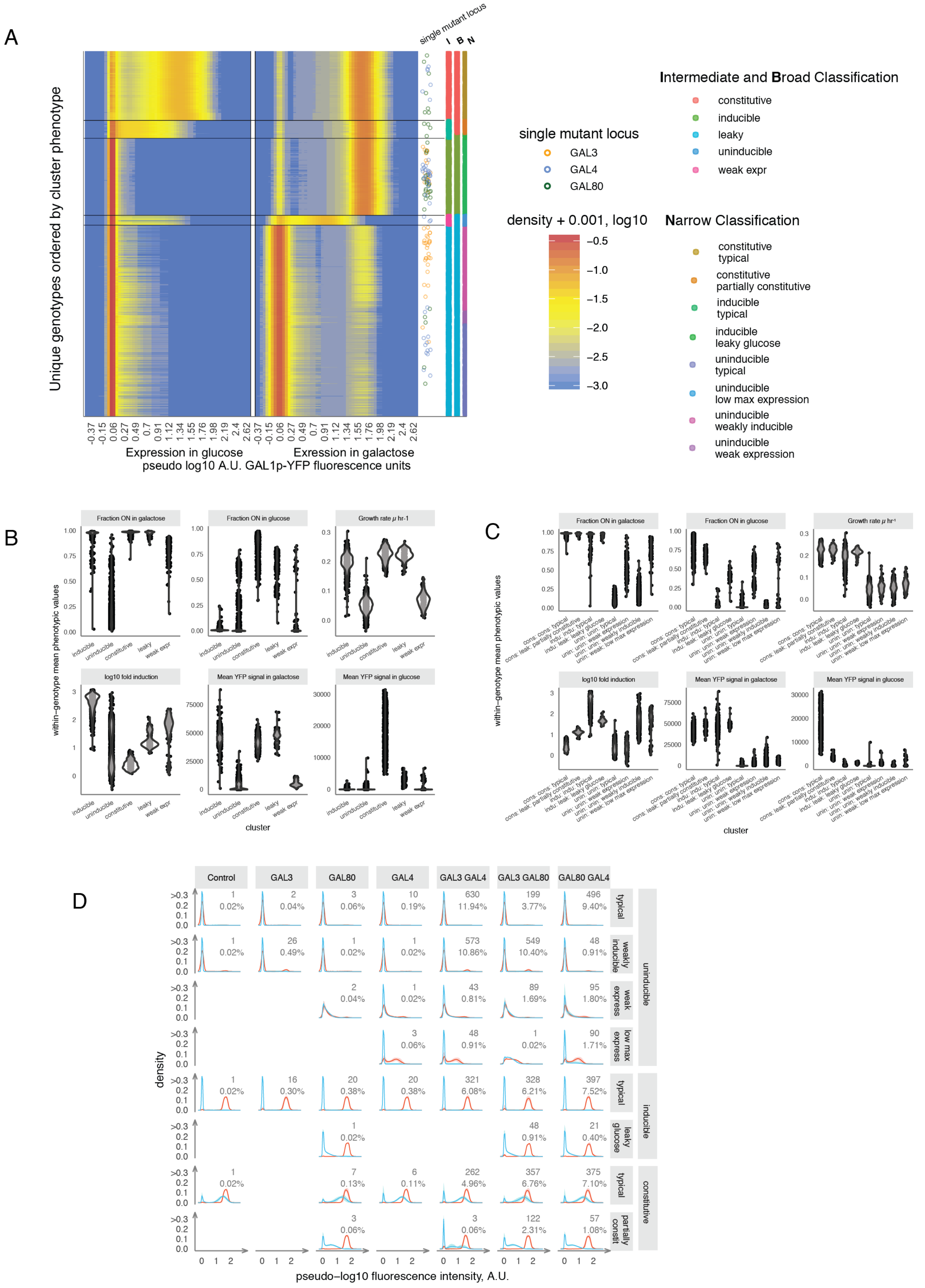

**Supplementary Figure 3: Distribution of phenotypes across classes, further subdivisions of gene expression distribution classes.**

All data presented in this figure arises from the second pairwise experiment. a) Transformed expression data used for clustering by the HDBScan* algorithm (Methods). The matrix used for clustering was of dimensions genotype (rows) x 120 A.U. fluorescence bins (cols). Each genotype's mean log10 density value + 0.001 for each A.U. fluorescence bin across in the glucose and galactose environments were used as values for clustering, and are coloured according to these values in the tile graph. The samples falling to the right and left of the white line delineates the measurements in glucose and galactose, respectively. The ordering of genotypes along the vertical axis was determined by mean within-cluster growth rate in galactose. The log of density values was used to exaggerate small differences for detection by the algorithm, e.g. the weak signal around leaky clones' expression in glucose. The "single mutant locus" points indicate where single mutants fell along the spectrum of expression profiles (vertical dimension, with horizontal jitter added to differentiate overlapping points), and are coloured according to which locus was mutated. The five classes reported in the text fell in the Intermediate set of classifications of a three-level hierarchy of classes (rightmost coloured lines, Methods). Most of these classifications arose "out of the box" depending on the cluster size setting used with the HDBScan* algorithm, however certain genotypes falling into infrequently observed or intermediate classes were assigned to clusters by hand (Methods). The rightmost three vertical lines are coloured corresponding to final classifications assigned to each clone for Intermediate, Broad and Narrow classifications. Black lines highlight transitions between intermediate classification groups.

b) and c): The distribution of phenotypic values across classes are shown as violin plots for the five key phenotypes, as well as the 'fold induction' measure used to sub-classify clones after initial unbiased clustering (Methods). Panel b) is how these characteristics broke down by intermediate classification, and c) is the same data broken down by narrow category. In c) the category labels follow class hierarchies as the broadest: intermediate: narrowest. d) the broadest and narrowest categories are plotted as in Fig 2C across controls, single and double mutants.

### Supplementary Figure 4

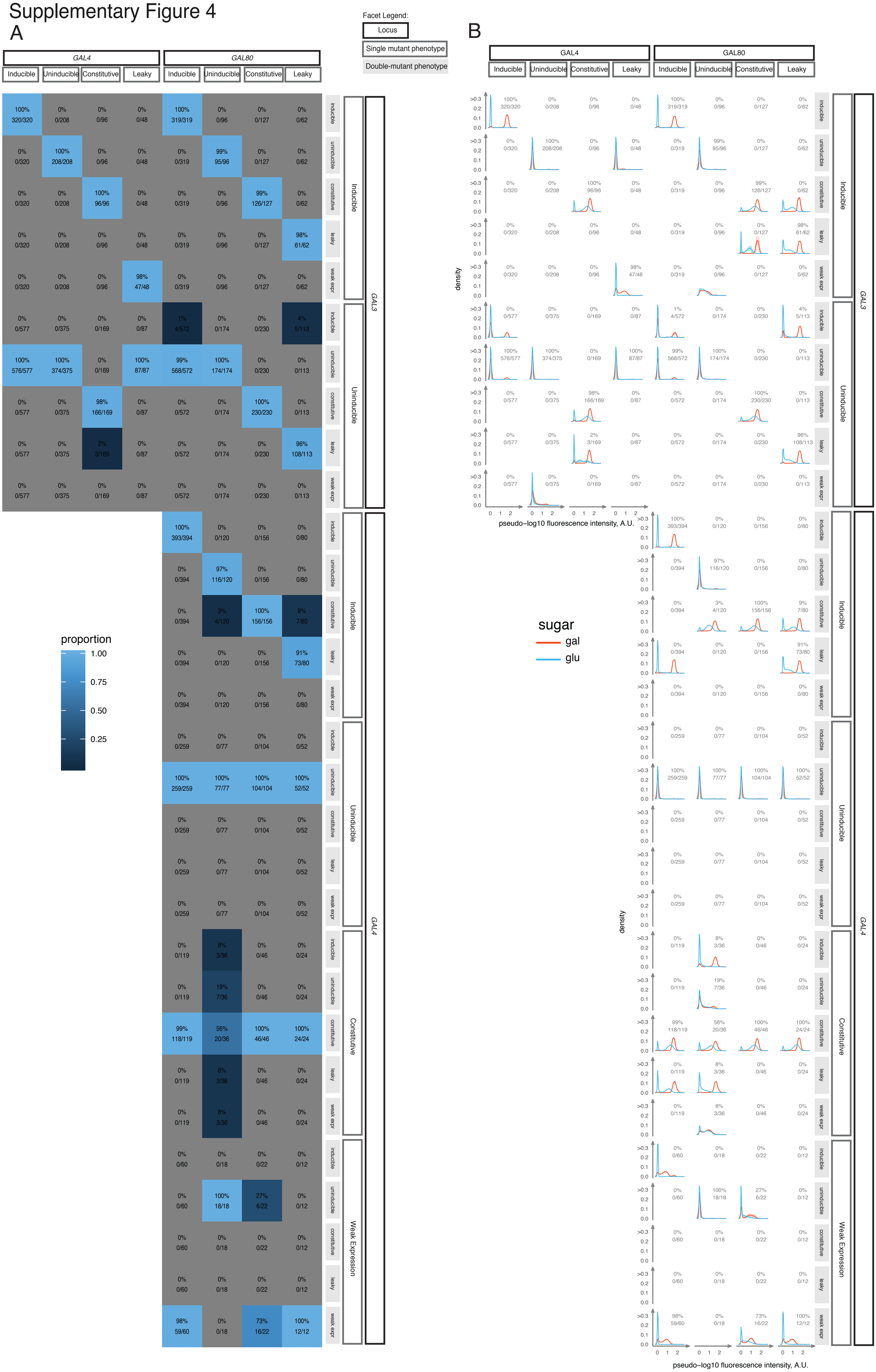

**Supplementary Figure 4: Double-mutant phenotypic classes arising from single-mutant class pairings.**

These plots illustrate the double mutant phenotypic classes which resulted from single mutants. The outermost facetting dimension encapsulates the locus, while the single-mutant class within locus is indicated in a second dimension. The vertical facetting includes a third dimension corresponding to the double-mutant phenotype that arose from the given pairing. a) Proportions of double mutant phenotype classes arising from the the given locus + single mutant pairing and b) Expression distribution histograms of all genotypes falling into the given locus + single-mutant + double-mutant expression class. Lines and shading are the mean and ± 1 SD of all genotypes falling into the given expression class. Blue lines are samples growing in glucose and red lines are samples after 12 hours of growth in galactose.

#### Supplementary Figure 5

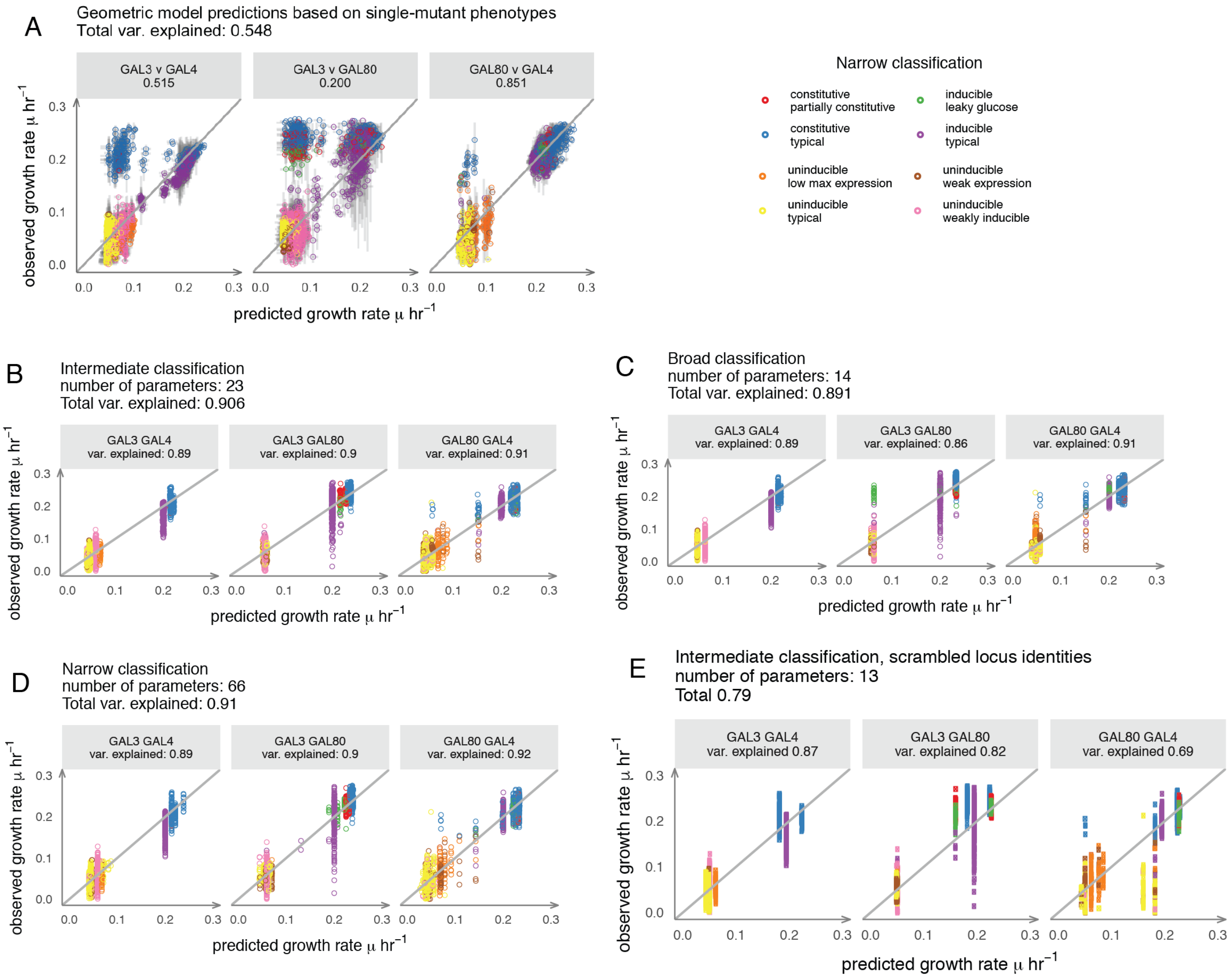

**Supplementary Figure 5: Predictability of growth rate based a geometric model of single-mutant effects or on single-mutant gene expression classes.**

The predicted vs observed values for prediction of double mutant phenotypes from a geometric model of growth rate variation based on single-mutant phenotypes (a) or allele phenotypes for classes from the three different hierarchies (b-e) (Methods). Datasets are facetted by the locus pairing and data points are coloured according to the Narrow classification scheme. In (a), as explained in the Main Text and Methods, the low variance explained stems from the fact that pathway-level epistasis leads to Constitutive and Leaky expression profiles (Fig 1D).

### Supplementary Figure 6

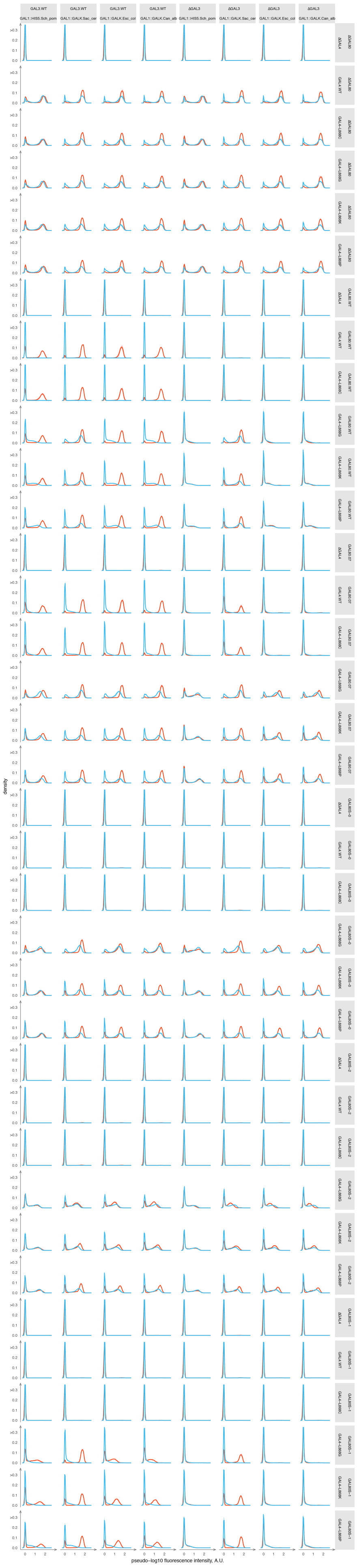

**Supplementary Figure 6: The distribution of *GAL1*pr-YFP expression across all single, double, triple and quadruple mutants from the third higher-order combinatorial mutant experiment.**

The genotypes of the GALR allele are indicated in the strip text at the top or right-hand side of each section's panel sets. Each panel is a unique genotype. Lines and shading are the mean and ± 1 SD of genotypes falling into the given genotype (N=4 for N=2 independent transformations). Blue lines are samples growing in glucose and red lines are samples after 12 hours of growth in galactose.

### Supplementary Figure 7

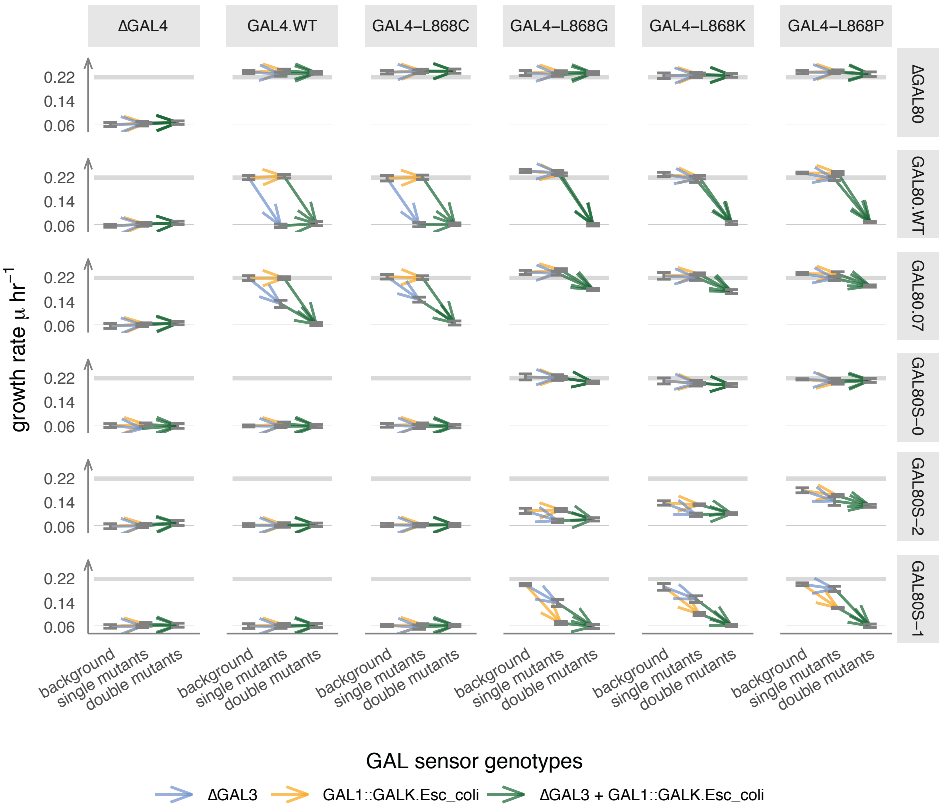

**Supplementary Figure 7: Complete landscape of the effects of *GAL1*::*GALK* and Δ*GAL3* on growth rate across *GAL4*-*GAL80* pairings**

Lines originate and terminate at mean values for the given genotype and grey bars indicate the 95% confidence interval.

#### Supplementary Figure 8

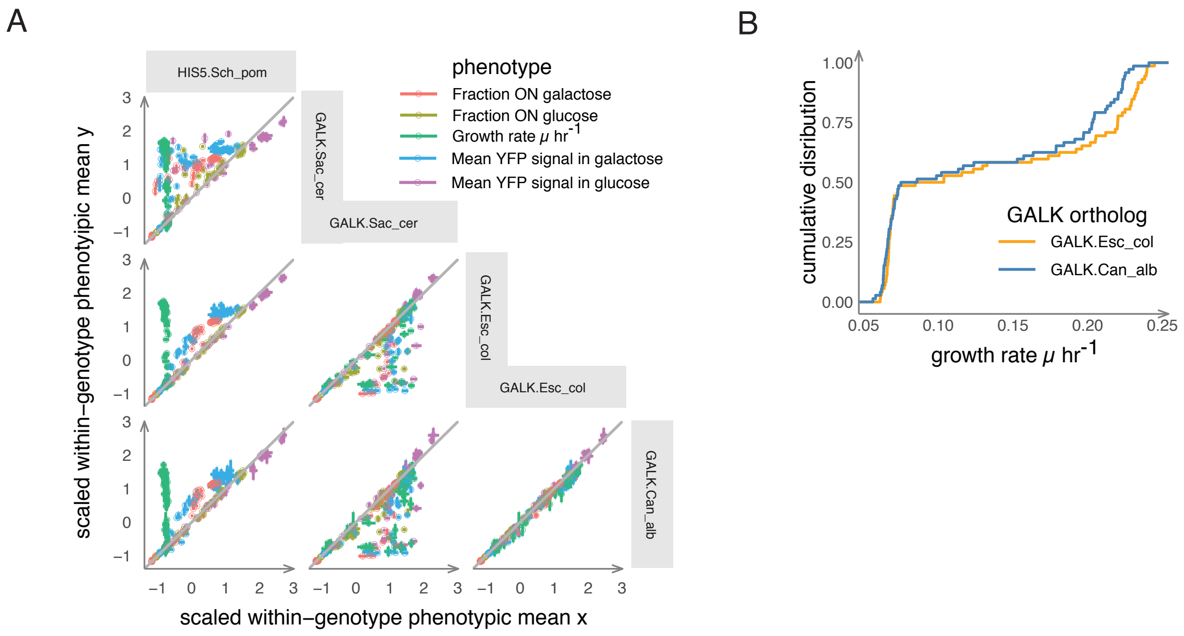

**Supplementary Figure 8: Pairwise comparisons of *GALK* backgrounds.**

All data presented in this figure arises from the third higher-order genetic interaction dataset. a) All within-genotype mean measurements were scaled (z-scores) within phenotypes. Each of the five phenotypes' mean scaled within-genotype values are plotted as points and error bars equal to 95% confidence interval. Each axis is a pairwise comparison between *GAL1pr-GALK* ortholog or the *GAL1pr-HIS5* from *S. pombe* constructs, with coloring of points and error bars corresponding to phenotypes. The width of error bars are equal to the 95% confidence interval. b) Phenotypes between *C. albicans* and *E. coli GALKs* were indistinguishable, except that *C. albicans* clones with high growth rates grew more slowly than genotype-matched *E. coli* clones (across individual observation, AD test statistic 5.273, $p < 0.002$, rejecting that samples come from the same distribution, whereas $p > 0.05$ for other phenotypes). Since *C. albicans* has a systematically lower growth rate, in plots and cartoons in figures 3B, 3D, 3E, and 4 we illustrate our data using backgrounds bearing *GAL1*::*GALK* from *E. coli*.

**Supplementary Table 1. Primers used in this study.**

| primer name | typical use | short description | sequence | amplifies | homology with | template | comments |
|---|---|---|---|---|---|---|---|
| AN087 | for cloning | AN087-GAL3-PromScan1-FP2 | CTAAACATAAAATCTGTAAAATAACAAG | GAL3 | NA | GAL3 | binds to ARS1 sequence upstream of GAL3; ~2500 bp product with AN089 |
| AN092 | for cloning | AN092-GAL3-CDSScan-reverse.primer2 | GCGGACATTTAGTCATATGTG | GAL3 | NA | GAL3 | binds 250 nt downstream of Y12 GAL3, 410 nt downstream of YPS/other alleles |
| AN112 | for cloning | AN112-GAL80amp[f]-plasmid | ATGGCGCAAGTTTTCCGCTTTGT | GAL80 | NA | GAL80 | for gibson cloning of GAL80 and GAL3 on the same plasmid. this primer amplifies gal80 with sticky bits homologous to plasmid makes ~2750 bp product |
| AN112 | for cloning | AN112-GAL80amp[f]-plasmid | ATGGCGCAAGTTTTCCGCTTTGT | GAL80 | cloning GAL80 | GAL80 | for gibson cloning of GAL80 and GAL3 on the same plasmid. this primer amplifies gal80 with sticky bits homologous to plasmid makes ~2750 bp product |
| AN153 | for cloning | AN153-GAL80-Scer-gibsoncloning-[r] | TTTATTTTTTGTGTTAGTGCACCTTCTC | GAL80 | NA | GAL80 | For cloning GAL80 using gibson cloning. To be used on genomic DNA template with primer AN109. generates 1828 bp fragment. |
| AN153 | for cloning | AN153-GAL80-Scer-gibsoncloning-[r] | TTTATTTTTTGTGTTAGTGCACCTTCTC | GAL80 | cloning GAL80 | GAL80 | For cloning GAL80 using gibson cloning. To be used on genomic DNA template with primer AN109. generates 1828 bp fragment. |
| ANgib133 | for cloning | pGAL4 | ttatacatcgttttgccttcttttaNNNNNNNNNNNNNgttggccgattcattaatgcag | pUC plasmid | GAL4 3' terminator | pUC19 | for cloning and GAL4 gene. |
| ANgib134 | for cloning | pGAL4 | gaataaaaaaaatactgggcgaatgctgtcgtttgcgtattgggcgctct | pUC plasmid | GAL4 5' promtoer | pUC19 | for cloning and GAL4 gene. |
| ANgib135 | for cloning | pGAL4 | gacagcattcgcccagtatt | GAL4 | plasmid reverse primer | GAL4 | GAL4_amp[f]. for cloning and library preparation of GAL4 genes |
| ANgib136 | for cloning | pGAL4 | taaaagaaggcaaaacgatgtataaat | GAL4 | plasmid forward primer | GAL4 | GAL4_amp[r]. for cloning and library preparation of GAL4 genes |
| ANgib158 | for cloning | GAL3_to_GAL4-[f] | atttatacatcgttttgccttcttttaGCCTAAACATAAAATCTGTAAAATAACAAG | GAL3 | GAL4 3' terminator | pGAL3 | to be used in joining PCR prods of GAL3 with 3' end of GAL4 |
| ANgib176 | for cloning | TEF.cassette.amp[r] | CAGTATAGCGACCAGCATTCA | TEFterm | NA | pRS series | generic for amplifying thigns with TEFterm |
| ANgib192 | for cloning | GAL3_to_GAL80 | ttcacgccagactatgtatgactcgatggcgcaagttttccgctttgt | GAL80 | GAL3 3' end | anything with S. cer GAL80 | for making fusions in plasmids with 3' end of GAL3. use with product generated when GAL3 amplified with AN110 as reverse primer. |
| ANgib196 | for cloning | GAL80.amp.NotI.pUC[r] | AAAGCTGGAGCTCCACCGCGGTGGCGGCCGCtttattttttgtgttagtgcaccttctc | GAL80 | NotI-digested pUC / pRS | GAL80 | for assembly of GAL genes into plasmids |
| ANgib197 | for cloning | GAL4.amp.NotI.pUC[f] | GGGATCCACTAGTTCTAGAGCGGCCGCgacagcattcgcccagtatt | GAL4 | NotI-digested pUC / pRS | GAL4 | for assembly of GAL genes into plasmids |
| ANgib201 | for cloning | GAL80_to_NotI_dig_pRS | GCTGGAGCTCCACCGCGGTGGCtttattttttgtgtcagtgcaccttctc | GAL80 | NotI-digested > Mungbeaned pRS plasmid | anything with S. cer GAL80 | for making plasmids for selecting GAL4 and GAL3 mutants. Use with ANgib192 as forward primer when making plasmid for GAL4 library. |
| ANgib202 | for cloning | GAL4_to_NotI_dig_pRS | GGGGATCCACTAGTTCTAGAGCGCgacagcattcgcccagtatt | GAL4 | NotI-digested > Mungbeaned pRS plasmid | GAL4 | for making plasmids for selecting GAL3 and GAL80 mutants. Use with ANgib158 as reverse primer for GAL80 mutant library. |
| ANgib203 | for cloning | GAL4.amp-NotI-Subcloneable | GGGGATCCACTAGTTCTAGAGCGCGTTTAAACTCTAACCAAATTAATCGAATTCTTGCGGCCGCgacagcattcgcccagtatt | GAL4 | 5' NotI digested BCinit | GAL4 | for use in generating the first plasmid for receiving the mutagenized libraries. NotI sites at each end, when digested, will yield sites ready for homologous recombination using the mutagenized and mutagenized PCR inserts. |
| ANgib204 | for cloning | GAL4.amp-NotI-Subcloneable | AAACGACGACAACTGGATGCGGCCGCtaaaagaaggcaaaacgatgtataaat | GAL4 | BC0 linker | GAL4 | for use in generating the first plasmid for receiving the mutagenized libraries. NotI sites at each end, when digested, will yield sites ready for homologous recombination using the mutagenized and mutagenized PCR inserts. |
| ANgib205 | for cloning | GAL3.BC0 linker | GGCCGCATCCAGTTGTCGTCGTTTAAACCTAAACATAAAATCTGTAAAATAACAAG | GAL3 | BC0 linker | GAL3 | for use in generating the first plasmid for receiving the mutagenized libraries. NotI sites at each end, when digested, will yield sites ready for homologous recombination using the mutagenized and mutagenized PCR inserts. |
| ANgib206 | for cloning | GAL4.BC1.5' linker | GCAGCAGATCGCCACATCCAGTTTAAACtaaaagaaggcaaaacgatgtataaat | GAL4 | BC1 linker | GAL4 | for use in generating the first plasmid for receiving the mutagenized libraries. NotI sites at each end, when digested, will yield sites ready for homologous recombination using the mutagenized and mutagenized PCR inserts. |
| ANgib207 | for cloning | GAL3.BC1.5' linker | AAACTGGATGTGGCGATCTGCTGCGGCCGCctaaacataaaatctgtaaaataacaag | GAL3 | BC1 linker | GAL3 | for use in generating the first plasmid for receiving the mutagenized libraries. NotI sites at each end, when digested, will yield sites ready for homologous recombination using the mutagenized and mutagenized PCR inserts. |
| ANgib208 | for cloning | GAL3.BC1.3' linker | AAACAGACGGCAGTCCTGCGGCGGCCGCcgagtcatacatagtctggcg | GAL3 | linkers | GAL3 | for use in generating the first plasmid for receiving the mutagenized libraries. NotI sites at each end, when digested, will yield sites ready for homologous recombination using the mutagenized and mutagenized PCR inserts. |
| ANgib209 | for cloning | GAL80.BC1.3'linker | GCCGCAGGACTGCCGTCTGTTTAAACatggcgcaagttttccgctttgt | GAL80 | linkers | GAL80 | for use in generating the first plasmid for receiving the mutagenized libraries. NotI sites at each end, when digested, will yield sites ready for homologous recombination using the mutagenized and mutagenized PCR inserts. |
| ANgib210 | for cloning | GAL3.BC2.5'linker | GCCGAAGCGGTGCTGGTGTCCGTTTAAACcgagtcatacatagtctggcg | GAL3 | linkers | GAL3 | for use in generating the first plasmid for receiving the mutagenized libraries. NotI sites at each end, when digested, will yield sites ready for homologous recombination using the mutagenized and mutagenized PCR inserts. |
| ANgib211 | for cloning | GAL80.BC2.5'linker | AAACGGACACCAGCACCGCTTCGGCGGCCGCatggcgcaagttttccgctttgt | GAL80 | linkers | GAL80 | for use in generating the first plasmid for receiving the mutagenized libraries. NotI sites at each end, when digested, will yield sites ready for homologous recombination using the mutagenized and mutagenized PCR inserts. |
| ANgib212 | for cloning | GAL80.BC2.3'linker | GCTGGAGCTCCACCGCGGTGGCGTTTAAACCTATATGCGTCACTGCCTGCGGCCGCtttatttttgtgtcagtgcaccttctc | GAL80 | NotI-digested > Mungbeaned pRS plasmid | GAL80 | for use in generating the first plasmid for receiving the mutagenized libraries. NotI sites at each end, when digested, will yield sites ready for homologous recombination using the mutagenized and mutagenized PCR inserts. |

| | | | | | | | |
|---|---|---|---|---|---|---|---|
| ANgib215 | for cloning | GAL4.libconstruct | TAACCAAATTAATCGAATTCTTGCgacagcattcgcccagtatt | GAL4 | 5' linker BC1 | GAL4 | for subcloning GAL4.S288c variants to assemble 3-gene screening plasmids |
| ANgib216 | for cloning | GAL4.libconstruct | GTTTAAACGACGACAACTGGATGCNNNNNNNNNAAACAGACGGCAGTCCTGCGGCGGCCGCAGCAGATCGCCACATCCAGTTTtaaaagaaggcaaaacgatgtataaat | GAL4 | 3' linker BC0 | GAL4 | for subcloning GAL4.S288c variants to assemble 3-gene screening plasmids |
| ANgib217 | for cloning | GAL3.libconstruct | AAACTGGATGTGGCGATCTGCTGCCTAAACATAAAATCTGTAAAATAACAAG | GAL3 | 5' linker BC2 | GAL3 | for subcloning GAL3.S288c variants to assemble 3-gene screening plasmids |
| ANgib218 | for cloning | GAL3.libconstruct | AAACAGACGGCAGTCCTGCGGCNNNNNNNNNAAACCTATATGCGTCACTGCCTGCGGCCGCCGAAGCGGTGCTGGTGTCCGTTTcgagtcatacatagtctggcg | GAL3 | 3' linker BC1 | GAL3 | for subcloning GAL3.S288c variants to assemble 3-gene screening plasmids |
| ANgib219 | for cloning | GAL80.libconstruct | AAACGGACACCAGCACCGCTTCGGCatggcgcaagttttccgctttgt | GAL80 | 5' linker BC3 | GAL80 | for subcloning GAL80.S288c variants to assemble 3-gene screening plasmids |
| ANgib220 | for cloning | GAL80.libconstruct | AAACCTATATGCGTCACTGCCTGCNNNNNNNNNAAACCAGCCCTGATACACTTGAGCGGCCGCtttattttttgtgtcagtgcaccttctc | GAL80 | 3' linker BC2 | GAL80 | for subcloning GAL80.S288c variants to assemble 3-gene screening plasmids |
| ANgib306 | for cloning | pRS413.amp.from.mcs.at.NotI | GCCACCGCGGTGGAGCTC | pRS series plasmids | MCS around NotI site | pRS plasmids | for amplifying the pRS plasmid for cloning |
| ANgib307 | for cloning | pRS413.amp.from.mcs.at.NotI | GCTCTAGAACTAGTGGATCC | pRS series plasmids | MCS around NotI site | pRS plasmids | for amplifying the pRS plasmid for cloning |
| ANgib320 | for cloning | GAL1pr-amp | tgctcattgctatattgaagtac | GAL1 promoter S288c | NA | S288c gDNA | NA |
| ANgib321 | for cloning | GAL1pr-YeCitrine-MET15 | GTATCGAAATGAGATGGCATggttgtttatgttcggatgt | TEFpr3' - whatever | homology with MET17/15 ORF. | pSR101 | homology with MET17/15 ORF. |
| ANgib322 | for cloning | MET 15 amp | ataaacaaccATGCCATCTCATTTCGATAC | MET17/15 | homology with TEF pr 3'. | BY4742 (LC50) | homology with TEF pr 3'. |
| ANgib349 | for cloning | plasmid amp f | GCTATACTGGTTTAAACCCTGCAGGcgctcacaattccacacaac | pRS series plasmids | TEF terminator | pRS series plasmids | introduces SbfI, PmeI sites |
| ANgib350 | for cloning | plasmid amp | cctatagtgagtcgtattacgc | pRS series plasmids | NA | pRS series plasmids | NA |
| ANgib351 | for cloning | URA3 frag amp | gcgtaatacgactcactataggCCTGCAGGGTTTAAACacagtcaacactaagacctatag | CaURA3 ORF 5' end from e.g. pSR101 | Vector reverse sequence (ANgib342 site) | pSR101 | NA |
| ANgib352 | for cloning | URA3 frag amp | ccgtacttcaatatagcaatgagcaGCGGCCGCGATGCACATAAATTGGTTTTCTTC | CaURA3 ORF 5' end from e.g. pSR101 | GAL1 promoter. | pSR101 | introduces NotI site |
| ANgib354 | for cloning | GAL1pr-amp | attgaactcaggtacaatcactt | GAL1 promoter S288c | NA | S288c gDNA | NA |
| ANgib355 | for cloning | E.coli.GALK.amp | AGaagtgattgtacctgagttcaatatgagtctgaaagaaaaaacacaa | GALK from E. coli | GAL1 promoter | E. coli gDNA | NA |
| ANgib356 | for cloning | E.coli.GALK.amp | TAAGAAATTCGCTTATTTAGAAGTGTTAGcactgtcctgctccttgtg | GALK from E. coli | ADH1 terminator | E. coli gDNA | NA |
| ANgib357 | for cloning | C. albicans GALK.amp | AGaagtgattgtacctgagttcaatatgtcagttcctacgtttgatg | GALK from C. albicans | GAL1 promoter | C. albicans gDNA | NA |
| ANgib358 | for cloning | C. albicans GALK.amp | TAAGAAATTCGCTTATTTAGAAGTGttattgtaagaatttcgtcgtt | GALK from C. albicans | ADH1 terminator | C. albicans gDNA | NA |
| ANgib359 | for cloning | HIS5.Spombe.amp | AGaagtgattgtacctgagttcaatATGGGTAGGAGGGCTTTTG | HIS5 from pUG27 (S. pombe derived) | GAL1 promoter | pUG27 | NA |
| ANgib360 | for cloning | HIS5.Spombe.amp | AAGAAATTCGCTTATTTAGAAGTGttacaacactcccttcgtgctt | HIS5 from pUG27 (S. pombe derived) | ADH1 terminator | pUG27 | NA |
| ANgib361 | for cloning | ADH.term.amp | CACTTCTAAATAAGCGAATTTCTTATG | ADH terminator and tef promoter from KT-series plasmid | NA | pKT-series plasmids | NA |
| ANgib362 | for cloning | ADH.term.amp emergency backup - if you can't amplify GALK's this will ligate with the GAL1 promoter fragment. Inserts FseI site flanked by 16 bp of random DNA | AGaagtgattgtacctgagttcaatGTTACAGGCCGGCCTACGCCACTTCTAAATAAGCGAATTTCTTATG | ADH terminator and tef promoter from KT-series plasmid | GAL1 promoter / first 15 aa of GAL1 | pKT-series plasmids | introduces FseI site and a bit of random DNA |
| ANgib363 | for cloning | MET 15 amp | TTATGGTTTTTGGCCAGCGAAA | MET17/15 | NA | BY4742 (LC50) | NA |

| | | | | | | | |
|---|---|---|---|---|---|---|---|
| ANgib364 | for cloning | TEF term amp | TGTTTTCGCTGGCCAAAAACCATAAGCGGCCGCTCAGTACTGACAATAAAAAGATTCT | TEF terminator | homology with MET15 CDS 3' | pSR101 | adds NotI site |
| ANgib365 | for cloning | TEF term amp | ttgtgagcgCCTGCAGGGTTTAAACCAGTATAGCGACCAGCATTCA | TEF terminator | homology with ANgib338 primer binding site | pSR101 | adds PmeI and SbfI sites |
| ANgib366 | for cloning | GAL1 ORF amp S288c | ATGACTAAATCTCATTCAGAAGAAG | GAL1 ORF from S288c | GAL1 promoter | S288c gDNA | NA |
| ANgib367 | for cloning | GAL1 ORF amp S288c | TAAGAAATTCGCTTATTTAGAAGTGTTATAATTCATATAGACAGCTGCC | GAL1 ORF from S288c | TEF terminator | S288c gDNA | NA |
| ANgib391 | for cloning | GAL4.locus.deletion 5' amp with GAL1pr binding site | AGGGGCGATTGGTTTGGGTGCGTGAGCGGCAAGAAGTTTCAAAACGTCCGCGTCCTTTGATgctcattgctatattgaagtac | pAMN45's GAL1pr-xx-Met15 cassette | GAL4 locus 5' region | pAMN45 - derived templates | for use with primer AN199 or ANgib176; integration into the GAL4 locus of strain AN634 |
| AN002 | for modifying yeast genome | GAL1_ProtFus_fp | GAGCTAGAAAATGCTATCATCGTCTCTAAACCAGCATTGGGCAGCTGTCTATATGAATTAatcggtgacggtgctggtttaatt | TEFpr-Selection Marker-TEFterm | 3' end GAL1 ORF | TEFpr-TEFterm e.g. pFA6 | for making in-frame fusions with GAL1 using pKT-series plasmids as template |
| AN003 | for modifying yeast genome | GAL1_FlPr_reverse.primer | aagttatgagtagaaaaaaatgagaagttgttctgaacaaagtaaaaaaaaagaagtatacgcataggccactagtggatctg | TEFpr-Selection Marker-TEFterm | 3' end of first 15 amino acids of GAL1 | TEFpr-TEFterm e.g. pFA6 | for making in-frame and promoter fusions with GAL1 using pKT-series plasmids as template |
| AN045 | for modifying yeast genome | GAL1 promoter fusion (f) | ggagaaaaaactataATGACTAAATCTCATTCAGAAGAAGTGATTGTACCTGAGTTCAATatcggtgacggtgctggtttaatt | TEFpr-Selection Marker-TEFterm | GAL1 terminator | TEFpr-TEFterm e.g. pFA6 | For use with pKT-series plasmids. For deleting GAL1 ORF. Keeps first 15 amino acids of GAL1. For use with GAL1_FlPr_reverse.primer |
| AN194 | for modifying yeast genome | AN194-ΔGAL80.locus[f] | GGTATTAACTTCTTCACTATAAGAAAATCACACGAGCGCCCGGACGATGTCTCTGTTTAACGACATGGAGGCCCAGAAT | TEFpr-Selection Marker-TEFterm | Upstream of GAL80 promoter | TEFpr-TEFterm e.g. pFA6 | for deleting the entire GAL80 locus (including promoer and terminator sequences) |
| AN195 | for modifying yeast genome | AN195-ΔGAL80.locus[r] | TTTGAAGCTATGATGGAAGGATGCCCGCTGCTGCAAAGTTTTGACAGTTATACGTAGCATCAGTATAGCGACCAGCATTCA | TEFpr-Selection Marker-TEFterm | Downstream of GAL80 terminaor | TEFpr-TEFterm e.g. pFA6 | for deleting the entire GAL80 locus (including promoer and terminator sequences) |
| AN196 | for modifying yeast genome | AN196-ΔGAL3.locus[f] | ttatttaagtattgtttgtgcacttgcctgcaggccttttgaaaagcaagcataaaagatCGACATGGAGGCCCAGAAT | TEFpr-Selection Marker-TEFterm | Upstream of GAL3 promoter | TEFpr-TEFterm e.g. pFA6 | for deleting the entire GAL3 locus (including promoer and terminator sequences) |
| AN197 | for modifying yeast genome | AN197-ΔGAL3.locus[r] | ttgcagggaatacaatggaaagacctaaagaaaatagaaaacggatttaaacggaaatggCAGTATAGCGACCAGCATTCA | TEFpr-Selection Marker-TEFterm | Downstream of GAL3 terminaor | TEFpr-TEFterm e.g. pFA6 | for deleting the entire GAL3 locus (including promoer and terminator sequences) |
| AN198 | for modifying yeast genome | AN198-ΔGAL4.locus[f] | AGGGGCGATTGGTTTGGGTGCGTGAGCGGCAAGAAGTTTCAAAACGTCCGCGTCCTTTGACGACATGGAGGCCCAGAAT | TEFpr-Selection Marker-TEFterm | Upstream of GAL4 promoter | TEFpr-TEFterm e.g. pFA6 | for deleting the entire GAL4 locus (including promoer and terminator sequences) |
| AN199 | for modifying yeast genome | AN199-ΔGAL4.locus[r] | TTCGTGAACTTCAGAGGCGATCGTATCTGTATTTTTCTCAATAGATTTGTCTGAAGACATCAGTATAGCGACCAGCATTCA | TEFpr-Selection Marker-TEFterm | Downstream of GAL4 terminaor | TEFpr-TEFterm e.g. pFA6 | for deleting the entire GAL4 locus (including promoer and terminator sequences) |
| ANgib391 | for modifying yeast genome | GAL4.locus.deletion 5' amp with GAL1pr binding site | AGGGGCGATTGGTTTGGGTGCGTGAGCGGCAAGAAGTTTCAAAACGTCCGCGTCCTTTGATgctcattgctatattgaagtac | TEFpr-Selection Marker-TEFterm | Upstream of GAL4 promoter | TEFpr-TEFterm e.g. pFA6 | for use with primer AN199 or ANgib176 on pAMN45 template to integrate constructs at GAL4::URA3 locus in strain AN634 |
| AN.TM.19 | targeted mutations | GAL3-ORF.deletion | AAGGTAGGGCAACACATAGTatgCACTAAACACCTTCTTGGAA | GAL3 | GAL3 | pAMN14 | complementary reverse primer is ANgib218 |
| AN.TM.21 | targeted mutations | GAL80-V352E | ATGATGCAGGTAAAGAAATCatggaagagTATCATTTACGAAATTATAATGCCATTG | GAL80 | GAL80 | pAMN15 | complementary reverse primer is ANgib220 |
| AN.TM.22 | targeted mutations | GAL80S-2 | ATGATGCAGGTAAAGAAATCatgaaggtaTATCATTTACGAAATTATAATGCCATTG | GAL80 | GAL80 | pAMN15 | complementary reverse primer is ANgib220 |
| AN.TM.25 | targeted mutations | GAL80S-1-G323R | ATCTGGTCCTTTACTACAGTcgtACTAGAGCAAACGACTTCCC | GAL80 | GAL80 | pAMN15 | complementary reverse primer is ANgib220 |

| | | | | | | | |
|---|---|---|---|---|---|---|---|
| AN.TM.26 | targeted mutations | GAL80S-0 | TTGACATTCACGGTACCAAGcgtGATTTGAAACTTGAAGGCGA | GAL80 | GAL80 | pAMN15 | complementary reverse primer is ANgib220 |
| AN.TM.34 | targeted mutations | GAL80-ORF.deletion | ccgttctttccactcccgtcATGaagcatcttgccctgtgctt | GAL80 | GAL80 | pAMN15 | complementary reverse primer is ANgib220 |
| AN.TM.36 | targeted mutations | GAL4-L868P | TGGATGATGgtaTATAACTATcccTTCGATGATGAAGATACCCC | GAL4 | GAL4 | pAMN31 | complementary reverse primer is ANgib216 |
| AN.TM.37 | targeted mutations | GAL4-L868X | TGGATGATGgtaTATAACTATnnsTTCGATGATGAAGATACCCC | GAL4 | GAL4 | pAMN31 | complementary reverse primer is ANgib216 |
| AN.TM.46 | targeted mutations | GAL4-ORF.deletion | agaagcaagcctcctgaaagATGaatgaatcgtagatactgaaaa | GAL4 | GAL4 | pAMN31 | complementary reverse primer is ANgib216 |
| AN.TM.61 | targeted mutations | GAL3-ORF.deletion | ACTATGTGTTGCCCTACCTT | GAL3 | GAL3 | pAMN14 | complementary reverse primer is ANgib217 |
| AN.TM.63 | targeted mutations | GAL80-V352E | GATTTCTTTACCTGCATCAT | GAL80 | GAL80 | pAMN15 | complementary reverse primer is ANgib219 |
| AN.TM.63 | targeted mutations | GAL80S-2 | GATTTCTTTACCTGCATCAT | GAL80 | GAL80 | pAMN15 | complementary reverse primer is ANgib219 |
| AN.TM.64 | targeted mutations | GAL80S-1-G323R | ACTGTAGTAAAGGACCAGAT | GAL80 | GAL80 | pAMN15 | complementary reverse primer is ANgib219 |
| AN.TM.65 | targeted mutations | GAL80S-0 | CTTGGTACCGTGAATGTCAA | GAL80 | GAL80 | pAMN15 | complementary reverse primer is ANgib219 |
| AN.TM.72 | targeted mutations | GAL80-ORF.deletion | gacgggagtggaaagaacgg | GAL80 | GAL80 | pAMN15 | complementary reverse primer is ANgib219 |
| AN.TM.74 | targeted mutations | GAL4-L868P | ATAGTTATAtacATCATCCATTG | GAL4 | GAL4 | pAMN31 | complementary reverse primer is ANgib215 |
| AN.TM.74 | targeted mutations | GAL4-L868X | ATAGTTATAtacATCATCCATTG | GAL4 | GAL4 | pAMN31 | complementary reverse primer is ANgib215 |
| AN.TM.80 | targeted mutations | GAL4-ORF.deletion | ctttcaggaggcttgcttct | GAL4 | GAL4 | pAMN31 | complementary reverse primer is ANgib215 |

**Supplementary Table 2. Strains and plasmids generated in this study.**

| yeast or bacteria | strain name | genotype | background | markers | yeast selection | parent strain | description and comments |
|---|---|---|---|---|---|---|---|
| yeast | KV447 | My copy of Kevin Verstrepen's copy of BY4741 | BY/S288c | :3Δ1 leu2Δ0 met15Δ | NA | BY4741 | [illegible] |
| yeast | SC5314 | Candida albicans standard lab strain from Tony Gabaldon's group | C. albicans | NA | NA | NA | [illegible] |
| yeast | AN566 | BY4741 GAL1-yeCitrine-KAN | BY/S288c | Kan+ | NA | KV447 | [illegible] |
| yeast | AN606 | AN566 ΔGAL80.locus::HYG | BY/S288c | HYG KAN | NA | AN566 | [illegible] |
| yeast | AN607 | AN566 ΔGAL3.locus::HYG | BY/S288c | HYG KAN | NA | AN566 | [illegible] |
| yeast | AN608 | AN566 ΔGAL4.locus::HYG | BY/S288c | HYG KAN | NA | AN566 | [illegible] |
| yeast | AN609 | AN606 (ΔGAL80.locus::HYG) ΔGAL4.locus::CaURA3 (from pSR101 template) | BY/S288c | HYG KAN CaURA3 | NA | AN606 | [illegible] |
| yeast | AN610 | AN607 (ΔGAL3.locus::HYG) ΔGAL4.locus::CaURA3 (from pSR101 template) | BY/S288c | HYG KAN CaURA3 | NA | AN607 | [illegible] |
| yeast | AN611 | AN607 (ΔGAL3.locus::HYG) ΔGAL80.locus::NAT | BY/S288c | HYG KAN NAT | NA | AN607 | [illegible] |
| yeast | AN612 | AN611 (ΔGAL3.locus::HYG) (ΔGAL80.locus::NAT) ΔGAL4.locus::CaURA3 (from pSR101 template) | BY/S288c | IYG KAN NAT CaURA | NA | AN611 | [illegible] |
| yeast | AN623 | KV447 ΔGAL1::YFP | BY/S288c | KAN | NA | KV447 | [illegible] |
| yeast | AN628 | AN623 ΔGAL3.locus::NAT | BY/S288c | KAN NAT | NA | AN623 | [illegible] |
| yeast | AN629 | AN628 (ΔGAL3.locus::NAT) ΔGAL80.locus::HYG (from pCB1 template) | BY/S288c | KAN NAT HYG | NA | AN628 | [illegible] |
| yeast | AN630 | AN623 ΔGAL4.locus::NAT | BY/S288c | KAN NAT | NA | AN623 | [illegible] |
| yeast | AN631 | AN623 ΔGAL80.locus::NAT | BY/S288c | KAN NAT | NA | AN623 | [illegible] |
| yeast | AN632 | AN630 (ΔGAL4.locus::NAT) ΔGAL3.locus::URA3 (from pSR101 template) | BY/S288c | KAN NAT URA3 | NA | AN630 | [illegible] |
| yeast | AN633 | AN631 (ΔGAL80.locus::NAT) ΔGAL4.locus::URA3 (from pSR101 template) | BY/S288c | KAN NAT URA3 | NA | AN631 | [illegible] |
| yeast | AN634 | AN629 (ΔGAL3.locus::NAT) (ΔGAL80.locus::HYG) ΔGAL4.locus::URA3 (from pSR101 template) | BY/S288c | KAN NAT HYG URA3 | NA | AN629 | [illegible] |
| yeast | AN636 | AN634 GALK.Ecoli AN634 transformed with linearized GALK construct from pAMN50.1 escherichia coli | BY/S288c | KAN NAT HYG MET | NA | AN634 | [illegible] |
| yeast | AN637 | AN634 GALK.Ecoli AN634 transformed with linearized GALK construct from pAMN50.2 escherichia coli | BY/S288c | KAN NAT HYG MET | NA | AN634 | [illegible] |
| yeast | AN638 | AN634 GALK.Can_alb AN634 transformed with linearized GALK construct from pAMN51.1 candida albicans | BY/S288c | KAN NAT HYG MET | NA | AN634 | [illegible] |
| yeast | AN639 | AN634 GALK.Sac_cer AN634 transformed with linearized GALK construct from pAMN52.2 saccharomyces cerevisiae | BY/S288c | KAN NAT HYG MET | NA | AN634 | [illegible] |
| yeast | AN640 | AN634 HIS5.Sch_pom AN634 transformed with linearized HIS5 construct from pAMN53.2 schizosaccharomyces pombe GAL1pr-his3 orthologue | BY/S288c | KAN NAT HYG MET | NA | AN634 | [illegible] |
| bacteria | pKT140 | From Kurt Thorne's plasmid collection (https://www.ncbi.nlm.nih.gov/pubmed/15197731) pKT-series plasmid with YeCitrine and KAN marker | pUC | AmpR | NA | NA | [illegible] |
| bacteria | pSR101 | pKT-series plasmid with mCherry cloned in place of fluorescent protein. Has CaURA3 cassette | pUC | AmpR | NA | NA | [illegible] |
| bacteria | pCB1 | A HYG-bearing pFA6-derived plasmid, originally from Brown, Murray, Verstrepen Current Biology 2009. | pUC | AmpR | NA | NA | [illegible] |
| bacteria | pAG25 | A NAT-bearing pFA6-derived plasmid | pUC | AmpR | NA | NA | [illegible] |
| bacteria | pUC19 | workhorse cloning vector | pUC | AmpR | NA | NA | [illegible] |
| bacteria | pUG27 | A HIS5-Sch_pombe-bearing pFA6-derived plasmid for complementing missing HIS3 activity in *S. cerevisiae* | pUC | AmpR | NA | NA | [illegible] |
| bacteria | pRS413 | yeast CEN plasmid with HIS3 selection | pUC | AmpR | HIS3 S. cer | NA | [illegible] |
| bacteria | pRS415 | yeast CEN plasmid with LEU2 selection | pUC | AmpR | LEU2 | NA | [illegible] |
| bacteria | pRS416 | yeast CEN plasmid with URA3 selection | pUC | AmpR | URA3 | NA | [illegible] |
| bacteria | pAMN14 | pGAL3 (pRS416 + GAL3 ORF) | pRS416 | AmpR | URA3 | pRS416 | [illegible] |
| bacteria | pAMN15 | pGAL80 (pRS416 + GAL80 ORF) | pRS416 | AmpR | URA3 | pRS416 | [illegible] |
| bacteria | pAMN26 | GAL4-GAL3-GAL80 GAL4 pre-screening plasmid. | pRS413 | AmpR | HIS3 S. cer | pRS413 | [illegible] |
| bacteria | pAMN27 | GAL4-GAL3-GAL80 GAL3 pre-screening plasmid. | pRS413 | AmpR | HIS3 S. cer | pRS413 | [illegible] |
| bacteria | pAMN28 | GAL4-GAL3-GAL80 GAL80 pre-screening plasmid. | pRS413 | AmpR | HIS3 S. cer | pRS413 | [illegible] |
| bacteria | pAMN31 | pGAL4 (pUC19 with GAL1) | pUC19 | AmpR | NA | pUC19 | [illegible] |
| bacteria | pAMN32 | pAMN26 ΔGAL4. NotI site for insertion of mutagenized GAL4 variants. | pRS413 / pAMN26 | AmpR | HIS3 S. cer | pAMN26 | [illegible] |
| bacteria | pAMN33 | pAMN27 ΔGAL3. NotI site for insertion of mutagenized GAL3 variants. | pRS413 / pAMN27 | AmpR | HIS3 S. cer | pAMN27 | [illegible] |
| bacteria | pAMN34 | pAMN28 ΔGAL80. NotI site for insertion of mutagenized GAL80 variants. | pRS413 / pAMN28 | AmpR | HIS3 S. cer | pAMN28 | [illegible] |
| bacteria | pAMN40 | GAL3 orf deletion | pRS413 / pAMN33 | AmpR | HIS3 S. cer | pAMN33 | [illegible] |
| bacteria | pAMN41 | GAL4 orf deletion | pRS413 / pAMN32 | AmpR | HIS3 S. cer | pAMN32 | [illegible] |
| bacteria | pAMN43 | GAL80 orf deletion | pRS413 / pAMN34 | AmpR | HIS3 S. cer | pAMN34 | [illegible] |
| bacteria | pAMN45 | pGAL1pr-xxx-MET15 -- see construction notes | pRS416 | AmpR | URA3 MET15 | pRS416 | [illegible] |
| bacteria | pAMN50 | pGAL1-Esc_col-GALK E. coli GALK | pRS416 / pAMN45 | AmpR | URA3 MET15 | bits n' pieces of pAMN45 | [illegible] |
| bacteria | pAMN51 | pGAL1-Can_alb-GALK candida albicans GALK | pRS416 / pAMN45 | AmpR | URA3 MET15 | bits n' pieces of pAMN45 | [illegible] |
| bacteria | pAMN52 | pGAL1-Sac_cer-GALK Saccharomyces cerevisiae control GAL1 | pRS416 / pAMN45 | AmpR | URA3 MET15 | bits n' pieces of pAMN45 | [illegible] |
| bacteria | pAMN53 | pGAL1-Sch_pom-HIS5 HIS3 complementation weird idea plasmid | pRS416 / pAMN45 | AmpR | URA3 MET15 HIS5 Sch. pom | bits n' pieces of pAMN45 | [illegible] |
| bacteria | GAL3.01 | GAL3.01 | pRS413 | GAL3.rand | HIS | pAMN33 | [illegible] |
| bacteria | GAL3.02 | GAL3.02 | pRS413 | GAL3.rand | HIS | pAMN33 | [illegible] |
| bacteria | GAL3.03 | GAL3.03 | pRS413 | GAL3.rand | HIS | pAMN33 | [illegible] |
| bacteria | GAL3.04 | GAL3.04 | pRS413 | GAL3.rand | HIS | pAMN33 | [illegible] |
| bacteria | GAL3.05 | GAL3.05 | pRS413 | GAL3.rand | HIS | pAMN33 | [illegible] |
| bacteria | GAL3.06 | GAL3.06 | pRS413 | GAL3.rand | HIS | pAMN33 | [illegible] |
| bacteria | GAL3.07 | GAL3.07 | pRS413 | GAL3.rand | HIS | pAMN33 | [illegible] |
| bacteria | GAL3.08 | GAL3.08 | pRS413 | GAL3.rand | HIS | pAMN33 | [illegible] |
| bacteria | GAL3.09 | GAL3.09 | pRS413 | GAL3.rand | HIS | pAMN33 | [illegible] |
| bacteria | GAL3.10 | GAL3.10 | pRS413 | GAL3.rand | HIS | pAMN33 | [illegible] |
| bacteria | GAL3.11 | GAL3.11 | pRS413 | GAL3.rand | HIS | pAMN33 | [illegible] |
| bacteria | GAL3.12 | GAL3.12 | pRS413 | GAL3.rand | HIS | pAMN33 | [illegible] |

| | | | | | | | |
|---|---|---|---|---|---|---|---|
| bacteria | GAL3.13 | GAL3.13 | pRS413 | GAL3.rand | HIS | pAMN33 | [illegible] |
| bacteria | GAL3.14 | GAL3.14 | pRS413 | GAL3.rand | HIS | pAMN33 | [illegible] |
| bacteria | GAL3.15 | GAL3.15 | pRS413 | GAL3.rand | HIS | pAMN33 | [illegible] |
| bacteria | GAL3.16 | GAL3.16 | pRS413 | GAL3.rand | HIS | pAMN33 | [illegible] |
| bacteria | GAL3.17 | GAL3.17 | pRS413 | GAL3.rand | HIS | pAMN33 | [illegible] |
| bacteria | GAL3.18 | GAL3.18 | pRS413 | GAL3.rand | HIS | pAMN33 | [illegible] |
| bacteria | GAL3.19 | GAL3.19 | pRS413 | GAL3.rand | HIS | pAMN33 | [illegible] |
| bacteria | GAL3.20 | GAL3.20 | pRS413 | GAL3.rand | HIS | pAMN33 | [illegible] |
| bacteria | GAL3.21 | GAL3.21 | pRS413 | GAL3.rand | HIS | pAMN33 | [illegible] |
| bacteria | GAL3.22 | GAL3.22 | pRS413 | GAL3.rand | HIS | pAMN33 | [illegible] |
| bacteria | GAL3.23 | GAL3.23 | pRS413 | GAL3.rand | HIS | pAMN33 | [illegible] |
| bacteria | GAL3.24 | GAL3.24 | pRS413 | GAL3.rand | HIS | pAMN33 | [illegible] |
| bacteria | GAL3.25 | GAL3.25 | pRS413 | GAL3.rand | HIS | pAMN33 | [illegible] |
| bacteria | GAL3.26 | GAL3.26 | pRS413 | GAL3.rand | HIS | pAMN33 | [illegible] |
| bacteria | GAL3.27 | GAL3.27 | pRS413 | GAL3.rand | HIS | pAMN33 | [illegible] |
| bacteria | GAL3.28 | GAL3.28 | pRS413 | GAL3.rand | HIS | pAMN33 | [illegible] |
| bacteria | GAL3.29 | GAL3.29 | pRS413 | GAL3.rand | HIS | pAMN33 | [illegible] |
| bacteria | GAL3.30 | GAL3.30 | pRS413 | GAL3.rand | HIS | pAMN33 | [illegible] |
| bacteria | GAL3.31 | GAL3.31 | pRS413 | GAL3.rand | HIS | pAMN33 | [illegible] |
| bacteria | GAL3.32 | GAL3.32 | pRS413 | GAL3.rand | HIS | pAMN33 | [illegible] |
| bacteria | GAL3.33 | GAL3.33 | pRS413 | GAL3.rand | HIS | pAMN33 | [illegible] |
| bacteria | GAL3.34 | GAL3.34 | pRS413 | GAL3.rand | HIS | pAMN33 | [illegible] |
| bacteria | GAL3.35 | GAL3.35 | pRS413 | GAL3.rand | HIS | pAMN33 | [illegible] |
| bacteria | GAL3.36 | GAL3.36 | pRS413 | GAL3.rand | HIS | pAMN33 | [illegible] |
| bacteria | GAL3.37 | GAL3.37 | pRS413 | GAL3.rand | HIS | pAMN33 | [illegible] |
| bacteria | GAL3.38 | GAL3.38 | pRS413 | GAL3.rand | HIS | pAMN33 | [illegible] |
| bacteria | GAL3.39 | GAL3.39 | pRS413 | GAL3.rand | HIS | pAMN33 | [illegible] |
| bacteria | GAL3.40 | GAL3.40 | pRS413 | GAL3.rand | HIS | pAMN33 | [illegible] |
| bacteria | GAL3.41 | GAL3.41 | pRS413 | GAL3.rand | HIS | pAMN33 | [illegible] |
| bacteria | GAL3.42 | GAL3.42 | pRS413 | GAL3.rand | HIS | pAMN33 | [illegible] |
| bacteria | GAL3.43 | GAL3.43 | pRS413 | GAL3.rand | HIS | pAMN33 | [illegible] |
| bacteria | GAL3.44 | GAL3.44 | pRS413 | GAL3.rand | HIS | pAMN33 | [illegible] |
| bacteria | GAL3.45 | GAL3.delta | pRS413 | GAL3.delta | HIS | pAMN33 | [illegible] |
| bacteria | GAL3.46 | GAL3.delta | pRS413 | GAL3.delta | HIS | pAMN33 | [illegible] |
| bacteria | GAL3.47 | GAL3.WT | pRS413 | GAL3.WT | HIS | pRS416 | [illegible] |
| bacteria | GAL3.48 | GAL3.WT | pRS413 | GAL3.WT | HIS | pRS416 | [illegible] |
| bacteria | GAL80.01 | GAL80.01 | pRS413 | GAL80.rand | HIS | pAMN34 | [illegible] |
| bacteria | GAL80.02 | GAL80.02 | pRS413 | GAL80.rand | HIS | pAMN34 | [illegible] |
| bacteria | GAL80.03 | GAL80.03 | pRS413 | GAL80.rand | HIS | pAMN34 | [illegible] |
| bacteria | GAL80.04 | GAL80.04 | pRS413 | GAL80.rand | HIS | pAMN34 | [illegible] |
| bacteria | GAL80.05 | GAL80.05 | pRS413 | GAL80.rand | HIS | pAMN34 | [illegible] |
| bacteria | GAL80.06 | GAL80.06 | pRS413 | GAL80.rand | HIS | pAMN34 | [illegible] |
| bacteria | GAL80.07 | GAL80.07 | pRS413 | GAL80.rand | HIS | pAMN34 | [illegible] |
| bacteria | GAL80.08 | GAL80.08 | pRS413 | GAL80.rand | HIS | pAMN34 | [illegible] |
| bacteria | GAL80.09 | GAL80.09 | pRS413 | GAL80.rand | HIS | pAMN34 | [illegible] |
| bacteria | GAL80.10 | GAL80.10 | pRS413 | GAL80.rand | HIS | pAMN34 | [illegible] |
| bacteria | GAL80.11 | GAL80.11 | pRS413 | GAL80.rand | HIS | pAMN34 | [illegible] |
| bacteria | GAL80.12 | GAL80.12 | pRS413 | GAL80.rand | HIS | pAMN34 | [illegible] |
| bacteria | GAL80.13 | GAL80.13 | pRS413 | GAL80.rand | HIS | pAMN34 | [illegible] |
| bacteria | GAL80.14 | GAL80.14 | pRS413 | GAL80.rand | HIS | pAMN34 | [illegible] |
| bacteria | GAL80.15 | GAL80.15 | pRS413 | GAL80.rand | HIS | pAMN34 | [illegible] |
| bacteria | GAL80.16 | GAL80.16 | pRS413 | GAL80.rand | HIS | pAMN34 | [illegible] |
| bacteria | GAL80.17 | GAL80.17 | pRS413 | GAL80.rand | HIS | pAMN34 | [illegible] |
| bacteria | GAL80.18 | GAL80.18 | pRS413 | GAL80.rand | HIS | pAMN34 | [illegible] |
| bacteria | GAL80.19 | GAL80.19 | pRS413 | GAL80.rand | HIS | pAMN34 | [illegible] |
| bacteria | GAL80.20 | GAL80.20 | pRS413 | GAL80.rand | HIS | pAMN34 | [illegible] |
| bacteria | GAL80.21 | GAL80.21 | pRS413 | GAL80.rand | HIS | pAMN34 | [illegible] |
| bacteria | GAL80.22 | GAL80.22 | pRS413 | GAL80.rand | HIS | pAMN34 | [illegible] |
| bacteria | GAL80.23 | GAL80.23 | pRS413 | GAL80.rand | HIS | pAMN34 | [illegible] |
| bacteria | GAL80.24 | GAL80.24 | pRS413 | GAL80.rand | HIS | pAMN34 | [illegible] |
| bacteria | GAL80.25 | GAL80.25 | pRS413 | GAL80.rand | HIS | pAMN34 | [illegible] |
| bacteria | GAL80.26 | GAL80.delta | pRS413 | GAL80.delta | HIS | pAMN34 | [illegible] |
| bacteria | GAL80.27 | GAL80.27 | pRS413 | GAL80.rand | HIS | pAMN34 | [illegible] |
| bacteria | GAL80.28 | GAL80.28 | pRS413 | GAL80.rand | HIS | pAMN34 | [illegible] |
| bacteria | GAL80.29 | GAL80.29 | pRS413 | GAL80.rand | HIS | pAMN34 | [illegible] |
| bacteria | GAL80.30 | GAL80.30 | pRS413 | GAL80.rand | HIS | pAMN34 | [illegible] |
| bacteria | GAL80.31 | GAL80S-V352E | pRS413 | V352E | HIS | pAMN34 | [illegible] |
| bacteria | GAL80.32 | GAL80S-V352E | pRS413 | V352E | HIS | pAMN34 | [illegible] |
| bacteria | GAL80.33 | GAL80.33 | pRS413 | GAL80.rand | HIS | pAMN34 | [illegible] |
| bacteria | GAL80.34 | GAL80.34 | pRS413 | GAL80.rand | HIS | pAMN34 | [illegible] |
| bacteria | GAL80.35 | GAL80S-2 | pRS413 | E351K | HIS | pAMN34 | [illegible] |
| bacteria | GAL80.36 | GAL80S-2 | pRS413 | E351K | HIS | pAMN34 | [illegible] |
| bacteria | GAL80.37 | GAL80S-1 | pRS413 | G323R | HIS | pAMN34 | [illegible] |
| bacteria | GAL80.38 | GAL80S-1 | pRS413 | G323R | HIS | pAMN34 | [illegible] |
| bacteria | GAL80.39 | GAL80S-1 | pRS413 | G323R | HIS | pAMN34 | [illegible] |
| bacteria | GAL80.40 | GAL80.40 | pRS413 | GAL80.rand | HIS | pAMN34 | [illegible] |
| bacteria | GAL80.41 | GAL80S-0 | pRS413 | G301R | HIS | pAMN34 | [illegible] |
| bacteria | GAL80.42 | GAL80S-0 | pRS413 | G301R | HIS | pAMN34 | [illegible] |
| bacteria | GAL80.43 | GAL80.43 | pRS413 | GAL80.rand | HIS | pAMN34 | [illegible] |
| bacteria | GAL80.44 | GAL80S-0 | pRS413 | G301R | HIS | pAMN34 | [illegible] |

| | | | | | | | |
|---|---|---|---|---|---|---|---|
| bacteria | GAL80.45 | GAL80.delta | pRS413 | GAL80.delta | HIS | pAMN34 | [illegible] |
| bacteria | GAL80.46 | GAL80.delta | pRS413 | GAL80.delta | HIS | pAMN34 | [illegible] |
| bacteria | GAL80.47 | GAL80.WT | pRS413 | GAL80.WT | HIS | pRS416 | [illegible] |
| bacteria | GAL80.48 | GAL80.WT | pRS413 | GAL80.WT | HIS | pRS416 | [illegible] |
| bacteria | GAL4.01 | GAL4.01 | pRS413 | GAL4.rand | HIS | pAMN32 | [illegible] |
| bacteria | GAL4.02 | GAL4.02 | pRS413 | GAL4.rand | HIS | pAMN32 | [illegible] |
| bacteria | GAL4.03 | GAL4.03 | pRS413 | GAL4.rand | HIS | pAMN32 | [illegible] |
| bacteria | GAL4.04 | GAL4.04 | pRS413 | GAL4.rand | HIS | pAMN32 | [illegible] |
| bacteria | GAL4.05 | GAL4.05 | pRS413 | GAL4.rand | HIS | pAMN32 | [illegible] |
| bacteria | GAL4.06 | GAL4.06 | pRS413 | GAL4.rand | HIS | pAMN32 | [illegible] |
| bacteria | GAL4.07 | GAL4.07 | pRS413 | GAL4.rand | HIS | pAMN32 | [illegible] |
| bacteria | GAL4.08 | GAL4.08 | pRS413 | GAL4.rand | HIS | pAMN32 | [illegible] |
| bacteria | GAL4.09 | GAL4.09 | pRS413 | GAL4.rand | HIS | pAMN32 | [illegible] |
| bacteria | GAL4.10 | GAL4.10 | pRS413 | GAL4.rand | HIS | pAMN32 | [illegible] |
| bacteria | GAL4.11 | GAL4.11 | pRS413 | GAL4.rand | HIS | pAMN32 | [illegible] |
| bacteria | GAL4.12 | GAL4.12 | pRS413 | GAL4.rand | HIS | pAMN32 | [illegible] |
| bacteria | GAL4.13 | GAL4.13 | pRS413 | GAL4.rand | HIS | pAMN32 | [illegible] |
| bacteria | GAL4.14 | GAL4.14 | pRS413 | GAL4.rand | HIS | pAMN32 | [illegible] |
| bacteria | GAL4.15 | GAL4.15 | pRS413 | GAL4.rand | HIS | pAMN32 | [illegible] |
| bacteria | GAL4.16 | GAL4.16 | pRS413 | GAL4.rand | HIS | pAMN32 | [illegible] |
| bacteria | GAL4.17 | GAL4.17 | pRS413 | GAL4.rand | HIS | pAMN32 | [illegible] |
| bacteria | GAL4.18 | GAL4.18 | pRS413 | GAL4.rand | HIS | pAMN32 | [illegible] |
| bacteria | GAL4.19 | GAL4.19 | pRS413 | GAL4.rand | HIS | pAMN32 | [illegible] |
| bacteria | GAL4.20 | GAL4.20 | pRS413 | GAL4.rand | HIS | pAMN32 | [illegible] |
| bacteria | GAL4.21 | GAL4.21 | pRS413 | GAL4.rand | HIS | pAMN32 | [illegible] |
| bacteria | GAL4.22 | GAL4.22 | pRS413 | GAL4.rand | HIS | pAMN32 | [illegible] |
| bacteria | GAL4.23 | GAL4.23 | pRS413 | GAL4.rand | HIS | pAMN32 | [illegible] |
| bacteria | GAL4.24 | GAL4.24 | pRS413 | GAL4.rand | HIS | pAMN32 | [illegible] |
| bacteria | GAL4.25 | GAL4.25 | pRS413 | GAL4.rand | HIS | pAMN32 | [illegible] |
| bacteria | GAL4.26 | GAL4.26 | pRS413 | GAL4.rand | HIS | pAMN32 | [illegible] |
| bacteria | GAL4.27 | GAL4.27 | pRS413 | GAL4.rand | HIS | pAMN32 | [illegible] |
| bacteria | GAL4.28 | GAL4.28 | pRS413 | GAL4.rand | HIS | pAMN32 | [illegible] |
| bacteria | GAL4.29 | GAL4.29 | pRS413 | GAL4.rand | HIS | pAMN32 | [illegible] |
| bacteria | GAL4.30 | GAL4.30 | pRS413 | GAL4.rand | HIS | pAMN32 | [illegible] |
| bacteria | GAL4.31 | GAL4.delta | pRS413 | GAL4.delta | HIS | pAMN32 | [illegible] |
| bacteria | GAL4.32 | GAL4.32 | pRS413 | GAL4.rand | HIS | pAMN32 | [illegible] |
| bacteria | GAL4.33 | GAL4.33 | pRS413 | GAL4.rand | HIS | pAMN32 | [illegible] |
| bacteria | GAL4.34 | GAL4.34 | pRS413 | GAL4.rand | HIS | pAMN32 | [illegible] |
| bacteria | GAL4.35 | GAL4-L868P | pRS413 | L868P | HIS | pAMN32 | [illegible] |
| bacteria | GAL4.36 | GAL4-L868C | pRS413 | L868C | HIS | pAMN32 | [illegible] |
| bacteria | GAL4.37 | GAL4-L868S | pRS413 | L868S | HIS | pAMN32 | [illegible] |
| bacteria | GAL4.38 | GAL4-L868G | pRS413 | L868G | HIS | pAMN32 | [illegible] |
| bacteria | GAL4.39 | GAL4-L868E | pRS413 | L868E | HIS | pAMN32 | [illegible] |
| bacteria | GAL4.40 | GAL4-L868K | pRS413 | L868K | HIS | pAMN32 | [illegible] |
| bacteria | GAL4.41 | GAL4-L868F | pRS413 | L868F | HIS | pAMN32 | [illegible] |
| bacteria | GAL4.42 | GAL4-L868S | pRS413 | L868S | HIS | pAMN32 | [illegible] |
| bacteria | GAL4.43 | GAL4-L868* | pRS413 | L868* | HIS | pAMN32 | [illegible] |
| bacteria | GAL4.44 | GAL4-L868G | pRS413 | L868G | HIS | pAMN32 | [illegible] |
| bacteria | GAL4.45 | GAL4.delta | pRS413 | GAL4.delta | HIS | pAMN32 | [illegible] |
| bacteria | GAL4.46 | GAL4.delta | pRS413 | GAL4.delta | HIS | pAMN32 | [illegible] |
| bacteria | GAL4.47 | GAL4.WT | pRS413 | GAL4.WT | HIS | pAMN32 | [illegible] |
| bacteria | GAL4.48 | GAL4.WT | pRS413 | GAL4.WT | HIS | pUC19 | [illegible] |

These materials were used for downstream combinatorial genetics experiments. Genotypes and associated phenotypes of the thousands of strains formed by combinatorial genetics experiments are included in the supplementary datasets. Basic plasmids all cloned / borne in DH5-alpha or its commercial derivative NEB 10-Beta

**Supplementary Table 3 - Generation of screening plasmids**

| primer(f) | primer(r) | amplifies | template | final.plasmid | locus | enzyme |
|---|---|---|---|---|---|---|
| ANgib203 | ANgib204 | GAL4 | pAMN31 | pAMN26 | GAL4 | ExTaq |
| ANgib205 | AN110 | GAL3 | pAMN14 | pAMN26 | GAL4 | ExTaq |
| ANgib192 | ANgib201 | GAL80 | pAMN15 | pAMN26 | GAL4 | ExTaq |
| ANgib202 | ANgib206 | GAL4 | pAMN31 | pAMN27 | GAL3 | ExTaq |
| ANgib207 | ANgib208 | GAL3 | pAMN14 | pAMN27 | GAL3 | ExTaq |
| ANgib209 | ANgib201 | GAL80 | pAMN15 | pAMN27 | GAL3 | ExTaq |
| ANgib202 | ANgib136 | GAL4 | pAMN31 | pAMN28 | GAL80 | ExTaq |
| ANgib158 | ANgib210 | GAL3 | pAMN14 | pAMN28 | GAL80 | ExTaq |
| ANgib211 | ANgib212 | GAL80 | pAMN15 | pAMN28 | GAL80 | ExTaq |

Co-transformation into strain AN612 of the three products from each group with NotI-digested pRS413 yielded plasmid assemblies that were transformed into *E. coli* and used downstream for cloning mutagenized alleles of GAL regulators. These plasmids are designed to have unique linker sequences flanking each gene that we are going to analyze by mutagenesis (locus.for.analysis column in the table).

**Supplementary Table 4 - Primers for random mutagenesis**

| primer.name | description | amplifies |
|---|---|---|
| ANgib215 | GAL4.libconstruct | GAL4 |
| ANgib216 | GAL4.libconstruct | GAL4 |
| ANgib217 | GAL3.libconstruct | GAL3 |
| ANgib218 | GAL3.libconstruct | GAL3 |
| ANgib219 | GAL80.libconstruct | GAL80 |
| ANgib220 | GAL80.libconstruct | GAL80 |

**Supplementary Table 5 - Primers for targeted mutagenesis**

| oligo.name | complementary primer | allele | fragment | enzyme |
|---|---|---|---|---|
| AN.TM.19 | ANgib218 | GAL3-ORF.deletion | 1 | ExTaq |
| AN.TM.21 | ANgib220 | GAL80-V352E | 1 | ExTaq |
| AN.TM.22 | ANgib220 | GAL80S-2 | 1 | ExTaq |
| AN.TM.25 | ANgib220 | GAL80S-1-G323R | 1 | ExTaq |
| AN.TM.26 | ANgib220 | GAL80S-0 | 1 | ExTaq |
| AN.TM.34 | ANgib220 | GAL80-ORF.deletion | 1 | ExTaq |
| AN.TM.36 | ANgib216 | GAL4-L868P | 1 | ExTaq |
| AN.TM.37 | ANgib216 | GAL4-L868X | 1 | ExTaq |
| AN.TM.46 | ANgib216 | GAL4-ORF.deletion | 1 | ExTaq |
| AN.TM.61 | ANgib217 | GAL3-ORF.deletion | 2 | ExTaq |
| AN.TM.63 | ANgib219 | GAL80-V352E | 2 | ExTaq |
| AN.TM.63 | ANgib219 | GAL80S-2 | 2 | ExTaq |
| AN.TM.64 | ANgib219 | GAL80S-1-G323R | 2 | ExTaq |
| AN.TM.65 | ANgib219 | GAL80S-0 | 2 | ExTaq |
| AN.TM.72 | ANgib219 | GAL80-ORF.deletion | 2 | ExTaq |
| AN.TM.74 | ANgib215 | GAL4-L868P | 2 | ExTaq |
| AN.TM.74 | ANgib215 | GAL4-L868X | 2 | ExTaq |
| AN.TM.80 | ANgib215 | GAL4-ORF.deletion | 2 | ExTaq |

All mutations were included in the fragment 1 product. Primers pairs above were used in separate reactions to generate the products for construction of targeted mutants.

**Supplementary Table 6 - PCR products for construction of targeted mutants**

| allele | oligo for fragment 1 | oligo for fragment 2 | plasmid | enzyme |
|---|---|---|---|---|
| GAL3-ORF.deletion | AN.TM.19 | AN.TM.61 | pAMN33 NotI and mungbeanned | ExTaq |
| GAL80-V352E | AN.TM.21 | AN.TM.63 | pAMN34 NotI and mungbeanned | ExTaq |
| GAL80S-2 | AN.TM.22 | AN.TM.63 | pAMN34 NotI and mungbeanned | ExTaq |
| GAL80S-1-G323R | AN.TM.25 | AN.TM.64 | pAMN34 NotI and mungbeanned | ExTaq |
| GAL80S-0 | AN.TM.26 | AN.TM.65 | pAMN34 NotI and mungbeanned | ExTaq |
| GAL80-ORF.deletion | AN.TM.34 | AN.TM.72 | pAMN34 NotI and mungbeanned | ExTaq |
| GAL4-L868P | AN.TM.36 | AN.TM.74 | pAMN32 NotI and mungbeanned | ExTaq |
| GAL4-L868X | AN.TM.37 | AN.TM.74 | pAMN32 NotI and mungbeanned | ExTaq |
| GAL4-ORF.deletion | AN.TM.46 | AN.TM.80 | pAMN32 NotI and mungbeanned | ExTaq |

**Supplementary Table 7 - PCR primers used in gap repair assembly for combinatorial genetics experiments**

| primer.name | locus | comments |
|---|---|---|
| ANgib197 | GAL4 alleles | 5' link with pRS-series plasmids amplified by ANgib306 x ANgib307 |
| ANgib136 | GAL4 alleles | 3' link with 5' end of GAL3 |
| AN110 | GAL3 alleles | 5' link with GAL4 3' end |
| ANgib158 | GAL3 alleles | 3' link with GAL80 5'end |
| ANgib192 | GAL80 alleles | 5' link with GAL3 3' end |

| | | |
|---|---|---|
| ANgib196 | GAL80 alleles | 3' link with pRS-series plasmids amplified by ANgib306 x ANgib307 |

These primers were used to generate products for assembly by gap repair

**Supplementary Table 8 - PCR reactions for combinatorial genetics experiments**

| rxn.num | primer(f) | primer(r) | template | amplifies | enzyme |
|---|---|---|---|---|---|
| 1 | AN110-GAL3amp[r]-GAL8( | ANgib158 | 14 or mutant-containing p | GAL3 alleles | KOD |
| 2 | ANgib192 | ANgib196 | 15 or mutant-containing p | GAL80 alleles | KOD |
| 3 | ANgib197 | ANgib136 | 31 or mutant-containing p | GAL4 alleles | KOD |
| 4 | ANgib306 | ANgib307 | pRS413 or pRS415 | vector LEU2 selection | Q5 |

All assemblies for combinatorial genetics used above 4 PCR products co-transformed into chassis strain AN612 or AN634.
pRS415 (LEU2 marker) was used for combinatorial genetics experiments.

**Supplementary Table 9 - PCR products used to generate pAMN45**

| rxn.num | primer(f) | primer(r) | template | amplifies | enzyme |
|---|---|---|---|---|---|
| 180209-prod.1 | ANgib349 | ANgib350 | pRS416 | vector | Q5 |
| 180209-prod.2 | ANgib351 | ANgib352 | pSR101 | cassette used to disrupt G | Q5 |
| 180209-prod.3 | ANgib320 | ANgib354 | S288c gDNA LC50 gDNA | GAL1 promoter | Q5 |
| 180209-prod.8 | ANgib362 | ANgib321 | pKT140 | and tef promoter from pK | Q5 |
| 180209-prod.9 | ANgib322 | ANgib363 | S288c gDNA LC50 gDNA | MET15 | Q5 |
| 180209-prod.10 | ANgib364 | ANgib365 | pSR101 | TEF terminator | Q5 |

These products were co-transformed into BY4741 and selected on SC-URA-MET plates. A single colony grew from this transformation and had mostly correct assembly junctions with a mutation in the PmeI restriction site.

**Supplementary Table 10 - PCR products used to generate GAL1pr-GALK constructs**

| rxn.num | primer(f) | primer(r) | template | amplifies | enzyme |
|---|---|---|---|---|---|
| 180213-prod3 | ANgib351 | ANgib354 | pAMN45 | 5' end construct | Q5 |
| 180213-prod4 | ANgib361 | ANgib365 | pAMN45 | 3' end construct | Q5 |
| 180213-prod5 | ANgib366 | ANgib367 | S288c gDNA LC50 gDNA | GAL1 | Q5 |
| 180209-prod.4 | ANgib355 | ANgib356 | E. coli (10^8 cells) | GALK E. coli | Q5 |
| 180209-prod.5 | ANgib357 | ANgib358 | C. albicans gDNA | GALK C. albicans | Q5 |
| 180209-prod.6 | ANgib359 | ANgib360 | pUG27 | HIS5 S. pombe | Q5 |

**Supplementary Table 11 - Transformations used to generate GAL1pr-GALK constructs**

| trans.num | prod1 | prod2 | prod3 | prod4 | construct | selection | final.plasmid |
|---|---|---|---|---|---|---|---|
| 180214-transformation1 | 180209-prod.1 | 180209-prod.4 | 180213-prod3 | 180213-prod4 | pGAL1pr-E.coli.GALK | -ura -met | pAMN50 |
| 180214-transformation2 | 180209-prod.1 | 180209-prod.5 | 180213-prod3 | 180213-prod4 | pGAL1pr-Calb.GALK | -ura -met | pAMN51 |
| 180214-transformation3 | 180209-prod.1 | 180209-prod.6 | 180213-prod3 | 180213-prod4 | pGAL1pr-HIS3.GALK | -ura -met | pAMN52 |
| 180214-transformation4 | 180209-prod.1 | 180213-prod5 (20 µl) | 180213-prod3 | 180213-prod4 | pGAL1pr-S.cer.GALK | -ura -met | pAMN53 |
