## Supplementary material for "Harmonious genetic combinations rewire regulatory networks and flip gene essentiality": Source data zip: Datasets-HeaderDescriptions.txt

variable	exp1	exp2	exp3	exvar	class	description
FITC.saturation.corrected.gal	exp1	exp2	exp3	0	logical	whether the sample had saturated FITC signal, in which case some values were inferred by PE-A signal
FITC.saturation.corrected.glu	exp1	exp2	exp3	0	logical	whether the sample had saturated FITC signal, in which case some values were inferred by PE-A signal
GAL3.WT	NA	NA	exp3	FALSE	logical	whether clone had this
GAL3.allele	NA	exp2	exp3	GAL3.delta	character	specific allele number (certain were duplicated within plates, each one getting a unique number)
GAL3.clone.named	NA	exp2	exp3	GAL3.delta	character	includes all replicates with the allele, such as "GAL3.WT" (sometimes different from the .allele)
GAL3.delta	NA	NA	exp3	TRUE	logical	whether clone had this
GAL3.mutant.class	NA	exp2	NA	rand	character	how the mutation was obatined
GAL4-L868C	NA	NA	exp3	FALSE	logical	whether clone had this
GAL4-L868G	NA	NA	exp3	FALSE	logical	whether clone had this
GAL4-L868K	NA	NA	exp3	FALSE	logical	whether clone had this
GAL4-L868P	NA	NA	exp3	FALSE	logical	whether clone had this
GAL4.WT	NA	NA	exp3	TRUE	logical	whether clone had this
GAL4.allele	NA	exp2	exp3	GAL4.WT	character	specific allele number (certain were duplicated within plates, each one getting a unique number)
GAL4.clone.named	NA	exp2	exp3	GAL4.WT	character	includes all replicates with the allele, such as "GAL3.WT" (sometimes different from the .allele)
GAL4.delta	NA	NA	exp3	FALSE	logical	whether clone had this
GAL4.mutant.class	NA	exp2	NA	WT	character	how the mutation was obatined
GAL80.07	NA	NA	exp3	FALSE	logical	whether clone had this
GAL80.WT	NA	NA	exp3	FALSE	logical	whether clone had this
GAL80.allele	NA	exp2	exp3	GAL80.37	character	specific allele number (certain were duplicated within plates, each one getting a unique number)
GAL80.clone.named	NA	exp2	exp3	GAL80S-1	character	includes all replicates with the allele, such as "GAL3.WT" (sometimes different from the .allele)
GAL80.delta	NA	NA	exp3	FALSE	logical	whether clone had this
GAL80.mutant.class	NA	exp2	NA	rand	character	how the mutation was obatined
GAL80S-0	NA	NA	exp3	FALSE	logical	whether clone had this
GAL80S-1	NA	NA	exp3	TRUE	logical	whether clone had this
GAL80S-2	NA	NA	exp3	FALSE	logical	whether clone had this
GALK.allele	NA	NA	exp3	pAMN50.1 - 2 - A2	character	specific allele number (certain were duplicated within plates, each one getting a unique number)
GALK.clone.named	NA	NA	exp3	GALK.Esc_col	character	includes all replicates with the allele, such as "GAL3.WT" (sometimes different from the .allele)
N.gal	exp1	exp2	exp3	5329	numeric	total number of events used to calculate statistics
N.glu	exp1	exp2	exp3	6439	numeric	total number of events used to calculate statistics
allele.named	NA	exp2	NA	GAL3.01 GAL80.01 GAL4.WT	character	the concatenated GALx.allele combination used for calculating statistics for unique genotypic combinations
arb	exp1	NA	NA	1	character	arbitrary factor used for pushing data around
background.strain	NA	exp2	NA	AN612	character	chassis strain used
binarycount	exp1	NA	NA	1 0 1	character	a genotype shorthand used in analyzing the first experiment
biol.rep	NA	NA	exp3	FALSE	character	the arbitrary nubmer of the biological rep
biol.rep.gal	NA	exp2	NA	1	factor	the arbitrary nubmer of the biological rep
biol.rep.glu	NA	exp2	NA	1	factor	the arbitrary nubmer of the biological rep
c	exp1	exp2	exp3	1	factor	column number in the 96-well plate
cell.per.ml.gal	exp1	NA	exp3	24531473.63	numeric	rates of events per ml from the cytometer calculated based on flow rate
cell.per.ml.glu	exp1	exp2	exp3	36394480.41	numeric	rates of events per ml from the cytometer calculated based on flow rate
cells.per.sec.gal	exp1	NA	exp3	12265.73681	numeric	rates of events from the cytometer
cells.per.sec.glu	exp1	NA	exp3	18197.2402	numeric	rates of events from the cytometer
clone.id	NA	NA	exp3	180324-Plate_001.A11	character	the date plate row column of the original transformant
clone.named	NA	exp2	NA	GAL3.01 GAL80.01 GAL4.WT	character	the concatenated GALx.clone.named combinations used for calculating statistics unique genotypic combinations
comments.glu	NA	exp2	NA	'NA'	character	comments on transformation or acquisition of the particular well
copies.GAL3.genome	exp1	NA	NA	1	numeric	number of copies of the gene in the genome
copies.GAL3.plasmid	exp1	NA	NA	0	numeric	number of copies of the gene in the plasmid
copies.GAL4.genome	exp1	NA	NA	1	numeric	number of copies of the gene in the genome
copies.GAL4.plasmid	exp1	NA	NA	0	numeric	number of copies of the gene in the plasmid
copies.GAL80.genome	exp1	NA	NA	0	numeric	number of copies of the gene in the genome
copies.GAL80.plasmid	exp1	NA	NA	0	numeric	number of copies of the gene in the plasmid
delta.GAL3	exp1	NA	NA	0	logical	whether clone had this
delta.GAL4	exp1	NA	NA	0	logical	whether clone had this
delta.GAL80	exp1	NA	NA	1	logical	whether clone had this
dens.-0.02.gal	exp1	NA	exp3	0	numeric	density header for AU fluorescence distributions
dens.-0.02.glu	exp1	NA	exp3	0.011	numeric	density header for AU fluorescence distributions
dens.-0.07.gal	exp1	NA	exp3	0	numeric	density header for AU fluorescence distributions
dens.-0.07.glu	exp1	NA	exp3	0.002	numeric	density header for AU fluorescence distributions
dens.-0.13.gal	exp1	NA	exp3	0	numeric	density header for AU fluorescence distributions
dens.-0.13.glu	exp1	NA	exp3	0	numeric	density header for AU fluorescence distributions
dens.-0.18.gal	exp1	NA	exp3	0	numeric	density header for AU fluorescence distributions
dens.-0.18.glu	exp1	NA	exp3	0	numeric	density header for AU fluorescence distributions
dens.-0.23.gal	exp1	NA	exp3	0	numeric	density header for AU fluorescence distributions
dens.-0.23.glu	exp1	NA	exp3	0	numeric	density header for AU fluorescence distributions
dens.-0.29.gal	exp1	NA	exp3	0	numeric	density header for AU fluorescence distributions
dens.-0.29.glu	exp1	NA	exp3	0	numeric	density header for AU fluorescence distributions
dens.-0.34.gal	exp1	NA	exp3	0	numeric	density header for AU fluorescence distributions
dens.-0.34.glu	exp1	NA	exp3	0	numeric	density header for AU fluorescence distributions
dens.-0.39.gal	exp1	NA	exp3	0	numeric	density header for AU fluorescence distributions
dens.-0.39.glu	exp1	NA	exp3	0	numeric	density header for AU fluorescence distributions
dens.-0.45.gal	exp1	NA	exp3	0	numeric	density header for AU fluorescence distributions
dens.-0.45.glu	exp1	NA	exp3	0	numeric	density header for AU fluorescence distributions
dens.-0.5.gal	exp1	NA	exp3	0	numeric	density header for AU fluorescence distributions
dens.-0.5.glu	exp1	NA	exp3	0	numeric	density header for AU fluorescence distributions
dens.0.03.gal	exp1	NA	exp3	0	numeric	density header for AU fluorescence distributions
dens.0.03.glu	exp1	NA	exp3	0.001	numeric	density header for AU fluorescence distributions
dens.0.09.gal	exp1	NA	exp3	0	numeric	density header for AU fluorescence distributions
dens.0.09.glu	exp1	NA	exp3	0	numeric	density header for AU fluorescence distributions
dens.0.14.gal	exp1	NA	exp3	0	numeric	density header for AU fluorescence distributions
dens.0.14.glu	exp1	NA	exp3	0	numeric	density header for AU fluorescence distributions
dens.0.19.gal	exp1	NA	exp3	0	numeric	density header for AU fluorescence distributions
dens.0.19.glu	exp1	NA	exp3	0	numeric	density header for AU fluorescence distributions
dens.0.25.gal	exp1	NA	exp3	0	numeric	density header for AU fluorescence distributions
dens.0.25.glu	exp1	NA	exp3	0	numeric	density header for AU fluorescence distributions
dens.0.3.gal	exp1	NA	exp3	0	numeric	density header for AU fluorescence distributions
dens.0.3.glu	exp1	NA	exp3	0	numeric	density header for AU fluorescence distributions
dens.0.35.gal	exp1	NA	exp3	0	numeric	density header for AU fluorescence distributions
dens.0.35.glu	exp1	NA	exp3	0	numeric	density header for AU fluorescence distributions
dens.0.41.gal	exp1	NA	exp3	0	numeric	density header for AU fluorescence distributions
dens.0.41.glu	exp1	NA	exp3	0	numeric	density header for AU fluorescence distributions
dens.0.46.gal	exp1	NA	exp3	0	numeric	density header for AU fluorescence distributions
dens.0.46.glu	exp1	NA	exp3	0	numeric	density header for AU fluorescence distributions
dens.0.51.gal	exp1	NA	exp3	0	numeric	density header for AU fluorescence distributions
dens.0.51.glu	exp1	NA	exp3	0	numeric	density header for AU fluorescence distributions
dens.0.57.gal	exp1	NA	exp3	0	numeric	density header for AU fluorescence distributions
dens.0.57.glu	exp1	NA	exp3	0	numeric	density header for AU fluorescence distributions
dens.0.62.gal	exp1	NA	exp3	0	numeric	density header for AU fluorescence distributions
dens.0.62.glu	exp1	NA	exp3	0	numeric	density header for AU fluorescence distributions
dens.0.67.gal	exp1	NA	exp3	0	numeric	density header for AU fluorescence distributions
dens.0.67.glu	exp1	NA	exp3	0.001	numeric	density header for AU fluorescence distributions
dens.0.73.gal	exp1	NA	exp3	0	numeric	density header for AU fluorescence distributions
dens.0.73.glu	exp1	NA	exp3	0.003	numeric	density header for AU fluorescence distributions
dens.0.78.gal	exp1	NA	exp3	0	numeric	density header for AU fluorescence distributions
dens.0.78.glu	exp1	NA	exp3	0.003	numeric	density header for AU fluorescence distributions
dens.0.83.gal	exp1	NA	exp3	0	numeric	density header for AU fluorescence distributions
dens.0.83.glu	exp1	NA	exp3	0.009	numeric	density header for AU fluorescence distributions
dens.0.89.gal	exp1	NA	exp3	0	numeric	density header for AU fluorescence distributions
dens.0.89.glu	exp1	NA	exp3	0.017	numeric	density header for AU fluorescence distributions
dens.0.94.gal	exp1	NA	exp3	0	numeric	density header for AU fluorescence distributions
dens.0.94.glu	exp1	NA	exp3	0.029	numeric	density header for AU fluorescence distributions
dens.0.99.gal	exp1	NA	exp3	0	numeric	density header for AU fluorescence distributions
dens.0.99.glu	exp1	NA	exp3	0.042	numeric	density header for AU fluorescence distributions
dens.1.05.gal	exp1	NA	exp3	0	numeric	density header for AU fluorescence distributions
dens.1.05.glu	exp1	NA	exp3	0.049	numeric	density header for AU fluorescence distributions
dens.1.1.gal	exp1	NA	exp3	0	numeric	density header for AU fluorescence distributions
dens.1.1.glu	exp1	NA	exp3	0.059	numeric	density header for AU fluorescence distributions
dens.1.15.gal	exp1	NA	exp3	0.001	numeric	density header for AU fluorescence distributions
dens.1.15.glu	exp1	NA	exp3	0.069	numeric	density header for AU fluorescence distributions
dens.1.21.gal	exp1	NA	exp3	0.005	numeric	density header for AU fluorescence distributions
dens.1.21.glu	exp1	NA	exp3	0.075	numeric	density header for AU fluorescence distributions
dens.1.26.gal	exp1	NA	exp3	0.022	numeric	density header for AU fluorescence distributions
dens.1.26.glu	exp1	NA	exp3	0.086	numeric	density header for AU fluorescence distributions
dens.1.31.gal	exp1	NA	exp3	0.05	numeric	density header for AU fluorescence distributions
dens.1.31.glu	exp1	NA	exp3	0.088	numeric	density header for AU fluorescence distributions
dens.1.37.gal	exp1	NA	exp3	0.094	numeric	density header for AU fluorescence distributions
dens.1.37.glu	exp1	NA	exp3	0.09	numeric	density header for AU fluorescence distributions
dens.1.42.gal	exp1	NA	exp3	0.143	numeric	density header for AU fluorescence distributions
dens.1.42.glu	exp1	NA	exp3	0.086	numeric	density header for AU fluorescence distributions
dens.1.47.gal	exp1	NA	exp3	0.174	numeric	density header for AU fluorescence distributions
dens.1.47.glu	exp1	NA	exp3	0.073	numeric	density header for AU fluorescence distributions
dens.1.53.gal	exp1	NA	exp3	0.177	numeric	density header for AU fluorescence distributions
dens.1.53.glu	exp1	NA	exp3	0.063	numeric	density header for AU fluorescence distributions
dens.1.58.gal	exp1	NA	exp3	0.161	numeric	density header for AU fluorescence distributions
dens.1.58.glu	exp1	NA	exp3	0.053	numeric	density header for AU fluorescence distributions
dens.1.63.gal	exp1	NA	exp3	0.093	numeric	density header for AU fluorescence distributions
dens.1.63.glu	exp1	NA	exp3	0.036	numeric	density header for AU fluorescence distributions
dens.1.69.gal	exp1	NA	exp3	0.043	numeric	density header for AU fluorescence distributions
dens.1.69.glu	exp1	NA	exp3	0.024	numeric	density header for AU fluorescence distributions
dens.1.74.gal	exp1	NA	exp3	0.017	numeric	density header for AU fluorescence distributions
dens.1.74.glu	exp1	NA	exp3	0.014	numeric	density header for AU fluorescence distributions
dens.1.79.gal	exp1	NA	exp3	0.011	numeric	density header for AU fluorescence distributions
dens.1.79.glu	exp1	NA	exp3	0.008	numeric	density header for AU fluorescence distributions
dens.1.85.gal	exp1	NA	exp3	0.005	numeric	density header for AU fluorescence distributions
dens.1.85.glu	exp1	NA	exp3	0.003	numeric	density header for AU fluorescence distributions
dens.1.9.gal	exp1	NA	exp3	0.003	numeric	density header for AU fluorescence distributions
dens.1.9.glu	exp1	NA	exp3	0.002	numeric	density header for AU fluorescence distributions
dens.1.95.gal	exp1	NA	exp3	0.001	numeric	density header for AU fluorescence distributions
dens.1.95.glu	exp1	NA	exp3	0	numeric	density header for AU fluorescence distributions
dens.2.01.gal	exp1	NA	exp3	0	numeric	density header for AU fluorescence distributions
dens.2.01.glu	exp1	NA	exp3	0	numeric	density header for AU fluorescence distributions
dens.2.06.gal	exp1	NA	exp3	0	numeric	density header for AU fluorescence distributions
dens.2.06.glu	exp1	NA	exp3	0	numeric	density header for AU fluorescence distributions
dens.2.11.gal	exp1	NA	exp3	0	numeric	density header for AU fluorescence distributions
dens.2.11.glu	exp1	NA	exp3	0	numeric	density header for AU fluorescence distributions
dens.2.17.gal	exp1	NA	exp3	0	numeric	density header for AU fluorescence distributions
dens.2.17.glu	exp1	NA	exp3	0	numeric	density header for AU fluorescence distributions
dens.2.22.gal	exp1	NA	exp3	0	numeric	density header for AU fluorescence distributions
dens.2.22.glu	exp1	NA	exp3	0	numeric	density header for AU fluorescence distributions
dens.2.27.gal	exp1	NA	exp3	0	numeric	density header for AU fluorescence distributions
dens.2.27.glu	exp1	NA	exp3	0	numeric	density header for AU fluorescence distributions
dens.2.33.gal	exp1	NA	exp3	0	numeric	density header for AU fluorescence distributions
dens.2.33.glu	exp1	NA	exp3	0	numeric	density header for AU fluorescence distributions
dens.2.38.gal	exp1	NA	exp3	0	numeric	density header for AU fluorescence distributions
dens.2.38.glu	exp1	NA	exp3	0	numeric	density header for AU fluorescence distributions
dens.2.43.gal	exp1	NA	exp3	0	numeric	density header for AU fluorescence distributions
dens.2.43.glu	exp1	NA	exp3	0	numeric	density header for AU fluorescence distributions
dens.2.49.gal	exp1	NA	exp3	0	numeric	density header for AU fluorescence distributions
dens.2.49.glu	exp1	NA	exp3	0	numeric	density header for AU fluorescence distributions
dens.2.54.gal	exp1	NA	exp3	0	numeric	density header for AU fluorescence distributions
dens.2.54.glu	exp1	NA	exp3	0	numeric	density header for AU fluorescence distributions
dens.2.59.gal	exp1	NA	exp3	0	numeric	density header for AU fluorescence distributions
dens.2.59.glu	exp1	NA	exp3	0	numeric	density header for AU fluorescence distributions
dens.2.65.gal	exp1	NA	exp3	0	numeric	density header for AU fluorescence distributions
dens.2.65.glu	exp1	NA	exp3	0	numeric	density header for AU fluorescence distributions
dens__-0.04665.gal	NA	exp2	NA	0.001	numeric	density header for AU fluorescence distributions
dens__-0.04665.glu	NA	exp2	NA	0.005	numeric	density header for AU fluorescence distributions
dens__-0.10015.gal	NA	exp2	NA	0	numeric	density header for AU fluorescence distributions
dens__-0.10015.glu	NA	exp2	NA	0	numeric	density header for AU fluorescence distributions
dens__-0.1535.gal	NA	exp2	NA	0	numeric	density header for AU fluorescence distributions
dens__-0.1535.glu	NA	exp2	NA	0	numeric	density header for AU fluorescence distributions
dens__-0.2065.gal	NA	exp2	NA	0	numeric	density header for AU fluorescence distributions
dens__-0.2065.glu	NA	exp2	NA	0	numeric	density header for AU fluorescence distributions
dens__-0.26.gal	NA	exp2	NA	0	numeric	density header for AU fluorescence distributions
dens__-0.26.glu	NA	exp2	NA	0	numeric	density header for AU fluorescence distributions
dens__-0.3135.gal	NA	exp2	NA	0	numeric	density header for AU fluorescence distributions
dens__-0.3135.glu	NA	exp2	NA	0	numeric	density header for AU fluorescence distributions
dens__-0.3665.gal	NA	exp2	NA	0	numeric	density header for AU fluorescence distributions
dens__-0.3665.glu	NA	exp2	NA	0	numeric	density header for AU fluorescence distributions
dens__-0.42.gal	NA	exp2	NA	0	numeric	density header for AU fluorescence distributions
dens__-0.42.glu	NA	exp2	NA	0	numeric	density header for AU fluorescence distributions
dens__-0.4735.gal	NA	exp2	NA	0	numeric	density header for AU fluorescence distributions
dens__-0.4735.glu	NA	exp2	NA	0	numeric	density header for AU fluorescence distributions
dens__0.00665.gal	NA	exp2	NA	0.005	numeric	density header for AU fluorescence distributions
dens__0.00665.glu	NA	exp2	NA	0.037	numeric	density header for AU fluorescence distributions
dens__0.06.gal	NA	exp2	NA	0.006	numeric	density header for AU fluorescence distributions
dens__0.06.glu	NA	exp2	NA	0.03	numeric	density header for AU fluorescence distributions
dens__0.11335.gal	NA	exp2	NA	0.003	numeric	density header for AU fluorescence distributions
dens__0.11335.glu	NA	exp2	NA	0.014	numeric	density header for AU fluorescence distributions
dens__0.1665.gal	NA	exp2	NA	0	numeric	density header for AU fluorescence distributions
dens__0.1665.glu	NA	exp2	NA	0.008	numeric	density header for AU fluorescence distributions
dens__0.22.gal	NA	exp2	NA	0	numeric	density header for AU fluorescence distributions
dens__0.22.glu	NA	exp2	NA	0.004	numeric	density header for AU fluorescence distributions
dens__0.2735.gal	NA	exp2	NA	0	numeric	density header for AU fluorescence distributions
dens__0.2735.glu	NA	exp2	NA	0.003	numeric	density header for AU fluorescence distributions
dens__0.3265.gal	NA	exp2	NA	0	numeric	density header for AU fluorescence distributions
dens__0.3265.glu	NA	exp2	NA	0.002	numeric	density header for AU fluorescence distributions
dens__0.38.gal	NA	exp2	NA	0.001	numeric	density header for AU fluorescence distributions
dens__0.38.glu	NA	exp2	NA	0.002	numeric	density header for AU fluorescence distributions
dens__0.4335.gal	NA	exp2	NA	0.001	numeric	density header for AU fluorescence distributions
dens__0.4335.glu	NA	exp2	NA	0.001	numeric	density header for AU fluorescence distributions
dens__0.4865.gal	NA	exp2	NA	0.001	numeric	density header for AU fluorescence distributions
dens__0.4865.glu	NA	exp2	NA	0.002	numeric	density header for AU fluorescence distributions
dens__0.54.gal	NA	exp2	NA	0.002	numeric	density header for AU fluorescence distributions
dens__0.54.glu	NA	exp2	NA	0.001	numeric	density header for AU fluorescence distributions
dens__0.5935.gal	NA	exp2	NA	0.001	numeric	density header for AU fluorescence distributions
dens__0.5935.glu	NA	exp2	NA	0.001	numeric	density header for AU fluorescence distributions
dens__0.6465.gal	NA	exp2	NA	0.001	numeric	density header for AU fluorescence distributions
dens__0.6465.glu	NA	exp2	NA	0.001	numeric	density header for AU fluorescence distributions
dens__0.7.gal	NA	exp2	NA	0.002	numeric	density header for AU fluorescence distributions
dens__0.7.glu	NA	exp2	NA	0.001	numeric	density header for AU fluorescence distributions
dens__0.7535.gal	NA	exp2	NA	0.001	numeric	density header for AU fluorescence distributions
dens__0.7535.glu	NA	exp2	NA	0.003	numeric	density header for AU fluorescence distributions
dens__0.8065.gal	NA	exp2	NA	0.002	numeric	density header for AU fluorescence distributions
dens__0.8065.glu	NA	exp2	NA	0.004	numeric	density header for AU fluorescence distributions
dens__0.86.gal	NA	exp2	NA	0.003	numeric	density header for AU fluorescence distributions
dens__0.86.glu	NA	exp2	NA	0.004	numeric	density header for AU fluorescence distributions
dens__0.9135.gal	NA	exp2	NA	0.003	numeric	density header for AU fluorescence distributions
dens__0.9135.glu	NA	exp2	NA	0.008	numeric	density header for AU fluorescence distributions
dens__0.9665.gal	NA	exp2	NA	0.005	numeric	density header for AU fluorescence distributions
dens__0.9665.glu	NA	exp2	NA	0.011	numeric	density header for AU fluorescence distributions
dens__1.0215.gal	NA	exp2	NA	0.003	numeric	density header for AU fluorescence distributions
dens__1.0215.glu	NA	exp2	NA	0.017	numeric	density header for AU fluorescence distributions
dens__1.075.gal	NA	exp2	NA	0.004	numeric	density header for AU fluorescence distributions
dens__1.075.glu	NA	exp2	NA	0.024	numeric	density header for AU fluorescence distributions
dens__1.125.gal	NA	exp2	NA	0.004	numeric	density header for AU fluorescence distributions
dens__1.125.glu	NA	exp2	NA	0.035	numeric	density header for AU fluorescence distributions
dens__1.18.gal	NA	exp2	NA	0.003	numeric	density header for AU fluorescence distributions
dens__1.18.glu	NA	exp2	NA	0.039	numeric	density header for AU fluorescence distributions
dens__1.235.gal	NA	exp2	NA	0.004	numeric	density header for AU fluorescence distributions
dens__1.235.glu	NA	exp2	NA	0.052	numeric	density header for AU fluorescence distributions
dens__1.285.gal	NA	exp2	NA	0.006	numeric	density header for AU fluorescence distributions
dens__1.285.glu	NA	exp2	NA	0.064	numeric	density header for AU fluorescence distributions
dens__1.34.gal	NA	exp2	NA	0.014	numeric	density header for AU fluorescence distributions
dens__1.34.glu	NA	exp2	NA	0.07	numeric	density header for AU fluorescence distributions
dens__1.395.gal	NA	exp2	NA	0.052	numeric	density header for AU fluorescence distributions
dens__1.395.glu	NA	exp2	NA	0.074	numeric	density header for AU fluorescence distributions
dens__1.445.gal	NA	exp2	NA	0.086	numeric	density header for AU fluorescence distributions
dens__1.445.glu	NA	exp2	NA	0.081	numeric	density header for AU fluorescence distributions
dens__1.5.gal	NA	exp2	NA	0.115	numeric	density header for AU fluorescence distributions
dens__1.5.glu	NA	exp2	NA	0.083	numeric	density header for AU fluorescence distributions
dens__1.555.gal	NA	exp2	NA	0.129	numeric	density header for AU fluorescence distributions
dens__1.555.glu	NA	exp2	NA	0.071	numeric	density header for AU fluorescence distributions
dens__1.605.gal	NA	exp2	NA	0.138	numeric	density header for AU fluorescence distributions
dens__1.605.glu	NA	exp2	NA	0.063	numeric	density header for AU fluorescence distributions
dens__1.66.gal	NA	exp2	NA	0.142	numeric	density header for AU fluorescence distributions
dens__1.66.glu	NA	exp2	NA	0.051	numeric	density header for AU fluorescence distributions
dens__1.715.gal	NA	exp2	NA	0.122	numeric	density header for AU fluorescence distributions
dens__1.715.glu	NA	exp2	NA	0.045	numeric	density header for AU fluorescence distributions
dens__1.765.gal	NA	exp2	NA	0.073	numeric	density header for AU fluorescence distributions
dens__1.765.glu	NA	exp2	NA	0.034	numeric	density header for AU fluorescence distributions
dens__1.82.gal	NA	exp2	NA	0.03	numeric	density header for AU fluorescence distributions
dens__1.82.glu	NA	exp2	NA	0.024	numeric	density header for AU fluorescence distributions
dens__1.875.gal	NA	exp2	NA	0.014	numeric	density header for AU fluorescence distributions
dens__1.875.glu	NA	exp2	NA	0.015	numeric	density header for AU fluorescence distributions
dens__1.925.gal	NA	exp2	NA	0.011	numeric	density header for AU fluorescence distributions
dens__1.925.glu	NA	exp2	NA	0.009	numeric	density header for AU fluorescence distributions
dens__1.98.gal	NA	exp2	NA	0.005	numeric	density header for AU fluorescence distributions
dens__1.98.glu	NA	exp2	NA	0.003	numeric	density header for AU fluorescence distributions
dens__2.035.gal	NA	exp2	NA	0.005	numeric	density header for AU fluorescence distributions
dens__2.035.glu	NA	exp2	NA	0.001	numeric	density header for AU fluorescence distributions
dens__2.085.gal	NA	exp2	NA	0.002	numeric	density header for AU fluorescence distributions
dens__2.085.glu	NA	exp2	NA	0	numeric	density header for AU fluorescence distributions
dens__2.14.gal	NA	exp2	NA	0	numeric	density header for AU fluorescence distributions
dens__2.14.glu	NA	exp2	NA	0.001	numeric	density header for AU fluorescence distributions
dens__2.195.gal	NA	exp2	NA	0	numeric	density header for AU fluorescence distributions
dens__2.195.glu	NA	exp2	NA	0	numeric	density header for AU fluorescence distributions
dens__2.245.gal	NA	exp2	NA	0	numeric	density header for AU fluorescence distributions
dens__2.245.glu	NA	exp2	NA	0	numeric	density header for AU fluorescence distributions
dens__2.3.gal	NA	exp2	NA	0	numeric	density header for AU fluorescence distributions
dens__2.3.glu	NA	exp2	NA	0	numeric	density header for AU fluorescence distributions
dens__2.355.gal	NA	exp2	NA	0	numeric	density header for AU fluorescence distributions
dens__2.355.glu	NA	exp2	NA	0	numeric	density header for AU fluorescence distributions
dens__2.405.gal	NA	exp2	NA	0	numeric	density header for AU fluorescence distributions
dens__2.405.glu	NA	exp2	NA	0	numeric	density header for AU fluorescence distributions
dens__2.46.gal	NA	exp2	NA	0	numeric	density header for AU fluorescence distributions
dens__2.46.glu	NA	exp2	NA	0	numeric	density header for AU fluorescence distributions
dens__2.515.gal	NA	exp2	NA	0	numeric	density header for AU fluorescence distributions
dens__2.515.glu	NA	exp2	NA	0	numeric	density header for AU fluorescence distributions
dens__2.565.gal	NA	exp2	NA	0	numeric	density header for AU fluorescence distributions
dens__2.565.glu	NA	exp2	NA	0	numeric	density header for AU fluorescence distributions
dens__2.62.gal	NA	exp2	NA	0	numeric	density header for AU fluorescence distributions
dens__2.62.glu	NA	exp2	NA	0	numeric	density header for AU fluorescence distributions
dens__2.675.gal	NA	exp2	NA	0	numeric	density header for AU fluorescence distributions
dens__2.675.glu	NA	exp2	NA	0	numeric	density header for AU fluorescence distributions
dip.pval.gal	exp1	exp2	exp3	0.988329423	numeric	dip statistic measuring whether samples were bimodal
dip.pval.glu	exp1	exp2	exp3	0.998264364	numeric	dip statistic measuring whether samples were bimodal
dip.stat.gal	exp1	exp2	exp3	0.003366014	numeric	dip statistic measuring whether samples were bimodal
dip.stat.glu	exp1	exp2	exp3	0.002328415	numeric	dip statistic measuring whether samples were bimodal
fac	NA	NA	exp3	GALK.Esc_col+GAL3.delta	character	something used for pushing data around
fold.dilution.gal	NA	NA	exp3	0.045	numeric	the magnitude of dilution from glucose into galactose
fold.dilution.glu	NA	NA	exp3	1	numeric	NA - the magnitude of dilution  glucose to glucose if applicable (this was never applicable in these experiments)
fracon.gal	exp1	exp2	exp3	0.999812348	numeric	fraction of cells ON in galactose
fracon.glu	exp1	exp2	exp3	0.98540146	numeric	fraction of cells ON in glucose
gal	NA	NA	exp3	0.2	numeric	concentration of galactose
gene	NA	NA	exp3	GALK	character	something used for pushing data around
generations	exp1	exp2	exp3	3.489807922	numeric	determined by merging day 1 glucose measurements with day 2 galactose measurements and calculating generations based on flow rates per ml
genomecopies	exp1	NA	NA	1 0 1	character	a summary of the number of copies of GAL regulators in the genome. not applicable for analyses discussed in the manuscript.
genotype.plasmid	exp1	NA	NA	vector	character	description of the plasmid
genotype.strain	exp1	NA	NA	AN606	character	description of the strain's genotype
genotype.strain1	exp1	NA	NA	GAL80.delta	character	description of the strain's genotype
glu	NA	NA	exp3	0	logical	whether or not glucose was there
growth.rate	exp1	exp2	exp3	0.20157921	numeric	the growth rate of the cells from the well determined based on the log of the fold change divided by the time grown in galactose
id	exp1	exp2	exp3	1	character	a factor used for joining samples from day 1 glucose to day 2 galactose
log.protein.production	NA	NA	exp3	4.706968898	numeric	total protein produced. used in data exploration and outside the scope of discussion here.
mean.fsc.gal	exp1	exp2	exp3	3579.472227	numeric	mean fsc signal for cells recorded from the well
mean.fsc.glu	exp1	exp2	exp3	3479.528964	numeric	mean fsc signal for cells recorded from the well
mean.norm.fsc.gal	exp1	exp2	exp3	9.726684173	numeric	in analysis from the raw flow cytometry, this was mean of each cell's YFP signal divided by fsc. in principle this controls for variation in volume of cells, but it didn't affect any analyses presented here.
mean.norm.fsc.glu	exp1	exp2	exp3	7.169440702	numeric	in analysis from the raw flow cytometry, this was mean of each cell's YFP signal divided by fsc. in principle this controls for variation in volume of cells, but it didn't affect any analyses presented here.
mean.on.fraction.gal	exp1	exp2	exp3	34114.40849	numeric	mean signal of cells that surpassed the "ON" threshold
mean.on.fraction.glu	exp1	exp2	exp3	23715.59935	numeric	mean signal of cells that surpassed the "ON" threshold
mean.on.fraction.glu_old	NA	NA	exp3	908.7272478	numeric	experiment 3 had a systematic bias in glucose measurements at the start of the day that could be controlled for at the mean level. this is the original measurement of mean YFP signal
mean.ssc.gal	exp1	exp2	exp3	4997.00292	numeric	mean ssc signal for cells recorded from the well
mean.ssc.glu	exp1	exp2	exp3	4128.152032	numeric	mean ssc signal for cells recorded from the well
measurement.date.gal	exp1	exp2	exp3	170928	numeric	the date of measurement, treated as a numeric
measurement.date.glu	exp1	exp2	exp3	170927	numeric	the date of measurement, treated as a numeric
median.fsc.gal	exp1	exp2	exp3	3426.5	numeric	median fsc signal for cells recorded from the well
median.fsc.glu	exp1	exp2	exp3	3329	numeric	median fsc signal for cells recorded from the well
median.ssc.gal	exp1	exp2	exp3	4936.399902	numeric	median ssc signal for cells recorded from the well
median.ssc.glu	exp1	exp2	exp3	3941	numeric	median ssc signal for cells recorded from the well
mixing_speed	NA	NA	exp3	200	numeric	speed of mixing, which was not applicable for any experiments because no mixing was used
mixing_speed.glu	NA	exp2	NA	200	numeric	speed of mixing, which was not applicable for any experiments because no mixing was used
mixing_volume	NA	NA	exp3	50	numeric	speed of mixing, which was not applicable for any experiments because no mixing was used
mixing_volume.glu	NA	exp2	NA	50	numeric	speed of mixing, which was not applicable for any experiments because no mixing was used
next.day	exp1	NA	NA	170928	numeric	used for merging glucose samples with next-day galactose samples
next.day.gal	NA	NA	exp3	NA	numeric	used for merging glucose samples with next-day galactose samples
next.day.glu	NA	exp2	exp3	180405	numeric	used for merging glucose samples with next-day galactose samples
number_mixes	NA	NA	exp3	0	numeric	mixing was not used for any experiments
number_mixes.glu	NA	exp2	NA	0	numeric	mixing was not used for any experiments
pRS415	exp1	NA	NA	1	logical	whether or not clone bore this element
parental.txant	NA	NA	exp3	1	logical	parental strain that was transformed
pct.oversaturation.gal	exp1	exp2	exp3	0	numeric	when FITC signal was oversaturated, this is the proportion of events that were oversaturated
pct.oversaturation.glu	exp1	exp2	exp3	0	numeric	when FITC signal was oversaturated, this is the proportion of events that were oversaturated
phenotypic.index	exp1	NA	exp3	1.985213807	numeric	the sum of fracon.glu + fracon.gal for each measured well
plasgeno	exp1	NA	NA	0 0 0 1 0 1	character	a shorthand summary of the genotype from experiment 1
plasmidcopies	exp1	NA	NA	0 0 0	character	a shorthand summary of the plasmid genotype from experiment 1
plate.gal	exp1	exp2	NA	Plate_009	character	plate number for the galactose measurement
plate.glu	exp1	exp2	NA	Plate_001	character	plate number for the glucose measurement
plate.num	NA	exp2	exp3	Plate_001	character	plate number
r	exp1	exp2	exp3	A	character	row of well
rc	exp1	exp2	exp3	A1	character	row column of well
record_sample_volume	NA	NA	exp3	10	numeric	unknown
record_sample_volume.glu	NA	exp2	NA	10	numeric	unknown
samp.plate.gal	NA	NA	exp3	180405-Plate_001	character	used for merging samples
samp.plate.glu	NA	NA	exp3	180404-Plate_001	character	used for merging samples
samp.plate.rc	exp1	NA	NA	Plate_001.A1	character	used for merging samples
samp_id.gal	exp1	exp2	exp3	170928-Plate_009.A1	character	a unique identifier for the measured well
samp_id.glu	exp1	exp2	exp3	170927-Plate_001.A1	character	a unique identifier for the measured well
sample_flow_rate	exp1	exp2	NA	0.5	numeric	the flow rate obtained from the template .xls file
sample_flow_rate.gal	NA	NA	exp3	2	numeric	the flow rate obtained from the template .xls file
sample_flow_rate.glu	NA	NA	exp3	0.5	numeric	the flow rate obtained from the template .xls file
sample_volume	NA	exp2	exp3	10	numeric	volume of cells measured
source.plate	exp1	exp2	exp3	Plate_073	character	the plate where the clone was stored in the freezer
source.plate.1	NA	NA	exp3	Plate_001	character	the plate where the clone was stored in the freezer
source.plate.suffix	NA	NA	exp3	1	character	the plate where the clone was stored in the freezer
strain.arb	exp1	NA	NA	strain.1	character	arbitrary name for the strain
total.copies.GAL3	exp1	NA	NA	1	numeric	the total copies of the given allele in the genome or plasmid
total.copies.GAL4	exp1	NA	NA	1	numeric	the total copies of the given allele in the genome or plasmid
total.copies.GAL80	exp1	NA	NA	0	numeric	the total copies of the given allele in the genome or plasmid
totalcopies	exp1	NA	NA	1 0 1	character	the total copies of all alleles in the genome or plasmid
tube.name	NA	NA	exp3	A11	character	the tube name from the script, basically the row and column of the plate
tube.name.glu	NA	exp2	NA	A1	character	the tube name from the script, basically the row and column of the plate
tx.c	NA	NA	exp3	11	character	the transformation's column
tx.mix.arb	exp1	NA	NA	tx.mix.1	character	the PCR mixture used for the independent transformation
tx.plate	NA	NA	exp3	Plate_001	character	the plate of the transformation
tx.plate.suffix	NA	NA	exp3	1	character	the plate of the transformation
tx.r	NA	NA	exp3	A	character	the row of the transformation
tx.rc	NA	NA	exp3	A11	character	the well of the transformed clone
tx.rep	NA	NA	exp3	1	character	the arbitrary number corresponding to the transformation replication
var.norm.fsc.gal	exp1	exp2	exp3	4.584178428	numeric	the variance in FSC-normalized YFP signal
var.norm.fsc.glu	exp1	exp2	exp3	6.992717814	numeric	the variance in FSC-normalized YFP signal
var.on.fraction.gal	exp1	exp2	exp3	100279720.7	numeric	the variance of mean YFP signal in ON cells
var.on.fraction.glu	exp1	exp2	exp3	166274051.3	numeric	the variance of mean YFP signal in ON cells
wash_volume	NA	NA	exp3	400	numeric	volume used for washing at the cytometer
wash_volume.glu	NA	exp2	NA	200	numeric	volume used for washing at the cytometer
yfp.25th.gal	exp1	exp2	exp3	27183.79883	numeric	quantiles of yfp expression
yfp.25th.glu	exp1	exp2	exp3	13820.09961	numeric	quantiles of yfp expression
yfp.5th.gal	exp1	exp2	exp3	20842.0793	numeric	quantiles of yfp expression
yfp.5th.glu	exp1	exp2	exp3	7800.659717	numeric	quantiles of yfp expression
yfp.75th.gal	exp1	exp2	exp3	39190.19922	numeric	quantiles of yfp expression
yfp.75th.glu	exp1	exp2	exp3	30200.09961	numeric	quantiles of yfp expression
yfp.95th.gal	exp1	exp2	exp3	51156	numeric	quantiles of yfp expression
yfp.95th.glu	exp1	exp2	exp3	48527.21797	numeric	quantiles of yfp expression
yfp.gal	NA	NA	exp3	119.1059565	numeric	mean YFP expression
yfp.glu	NA	NA	exp3	8.989528553	numeric	mean YFP expression
yfp.max.gal	exp1	exp2	exp3	102859.3984	numeric	quantiles of yfp expression
yfp.max.glu	exp1	exp2	exp3	122773	numeric	quantiles of yfp expression
yfp.mean.gal	exp1	exp2	exp3	34108.04283	numeric	mean YFP expression
yfp.mean.glu	exp1	exp2	exp3	23369.32186	numeric	mean YFP expression
yfp.mean.glu_old	NA	NA	exp3	52.44888648	numeric	experiment 3 had a systematic bias in glucose measurements at the start of the day that could be controlled for at the mean level. this is the original measurement of mean YFP signal
yfp.median.gal	exp1	exp2	exp3	32912.59766	numeric	quantiles of yfp expression
yfp.median.glu	exp1	exp2	exp3	20860	numeric	quantiles of yfp expression
yfp.min.gal	exp1	exp2	exp3	191.8000031	numeric	quantiles of yfp expression
yfp.min.glu	exp1	exp2	exp3	-175	numeric	quantiles of yfp expression
yfp.variance.gal	exp1	exp2	exp3	100476839.3	numeric	total variance of YFP signal
yfp.variance.glu	exp1	exp2	exp3	171941409.5	numeric	total variance of YFP signal
